# Learning to learn persistently modifies an entorhinal-hippocampal excitatory-inhibitory subcircuit

**DOI:** 10.1101/817627

**Authors:** Ain Chung, Claudia Jou, Alejandro Grau-Perales, Eliott Levy, Dino Dvorak, Nida Hussain, André A. Fenton

**Affiliations:** Center for Neural Science, New York University; Department of Biological Sciences, City University of New York, Hunter College; Neuroscience Institute at the NYU Langone Medical Center

## Abstract

Cognitive control, the judicious use of relevant information while ignoring distractions, is a feature of everyday cognitive experience, but its neurobiology is understudied. We investigated whether cognitive control training (CCT) changes hippocampal neural circuit function in mice, beyond the changes caused by place learning and memory formation. Mice learned and remembered a conditioned place avoidance during CCT that required ignoring irrelevant locations of shock. They were compared to controls that learned the same place avoidance under lower cognitive control demands. Weeks after CCT, mice learn new tasks in novel environments faster than controls; they learned to learn. We investigated entorhinal cortex-to-dentate gyrus neural circuit changes and report that CCT rapidly changes synaptic circuit function, resulting in an excitatory-inhibitory subcircuit change that persists for months. CCT increases inhibition that attenuates the dentate response to medial entorhinal cortical input, and through disinhibition, potentiates the response to strong inputs, pointing to overall signal-to-noise enhancement. These neurobiological findings support a neuroplasticity hypothesis that, beyond storing item/event associations, CCT persistently optimizes neural circuit information processing.

---

Neurobiological investigations of learning typically focus on the acquisition of memory for content-specific information, or even generalization of that information across distinct contexts ^1,2^, whereas outside the laboratory, acquiring a particular memory often occurs concurrently with learning to learn ^1–4^. Learning to learn relies on acquiring cognitive skills to make decisions in the presence of noisy data ^5^, forming so-called schema, a cognitive framework for information processing that is abstracted from the particular content of memory ^6,7^ and can be the basis of a particular mindset that directs future learning and experience ^8^. Learning to learn is the basis of the cognitive behavioral therapist’s effort to improve subsequent understanding, learning and overall function of their patients ^9^. While it is commonplace to suppose that the cognitive psychology of schema, the psychotherapeutic practice of cognitive behavioral therapy (CBT), and the neurobiology of memory and neural information processing are intimately related, the nature of those relations are speculative ^9^. A commonly-stated neuroplasticity hypothesis asserts that cognitive training causes neural plasticity to change brain function ^10,11^, but beyond changes in sensory systems ^12,13^, there is scant evidence for the three key predictions underlying this hypothesis 1) cognitive training-induced 2) long-lasting, and 3) memory-nonspecific training-induced changes in neural circuit function.

Here, we test these predictions using freely-behaving mice. We exploited an experimental platform that demonstrated persistent *ex vivo* and anesthetized *in vivo* electrophysiological changes in GABAergic-sensitive hippocampus synaptic function after learning an active place avoidance task ^14^. The task requires intact hippocampal activity ^15,16^, and persistent PKMzeta (PKMζ)-mediated long-term potentiation (LTP) of hippocampal synaptic function ^17–20^.

We begin by demonstrating cognitive control of switching between task relevant and irrelevant distracting information in the activity of mouse hippocampus CA1 during the active place avoidance task variant in which mice on a rotating (1 rpm) arena learn the (“relevant”), room-based stationary location of a shock zone, but must not associate shock with (“distractions”), the unpredictable arena-based rotating locations where they experience shock ^21–23^. Hippocampus CA1 switches between representing room and arena locations during pretraining when the mouse is first exposed to the rotating arena environment without reinforcement (Fig. 1). However, during and after training to avoid shock reinforcement, CA1 activity alternates between encoding task-relevant room locations near the room-defined shock zone and task-irrelevant arena locations far from the shock zone (Fig. 1e), demonstrating representational multi-tasking that mirrors ongoing goal-directed performance, the defining feature of cognitive control ^21,22,24^.

**Figure 1:**
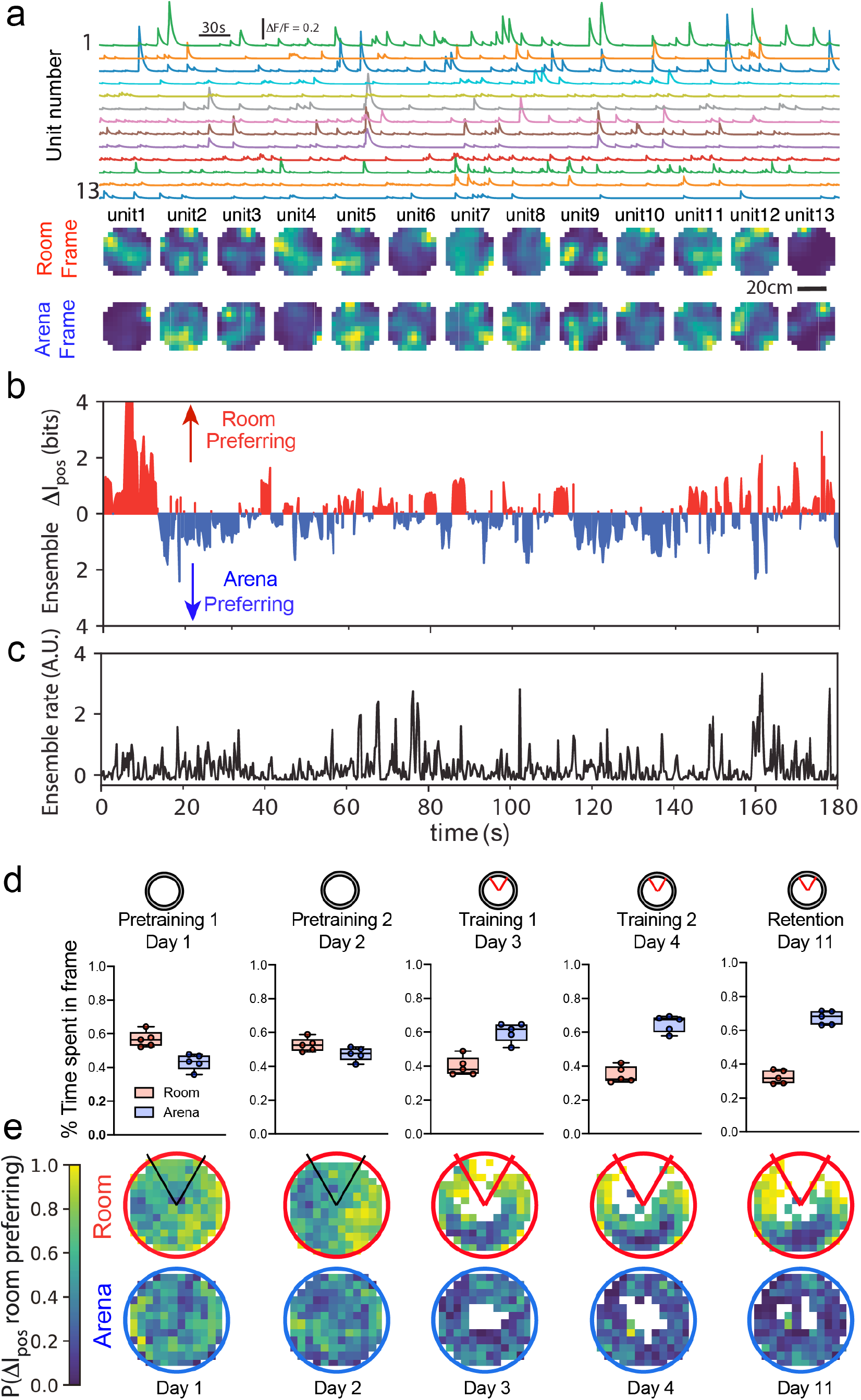
Cognitive control of spatial information processing in mouse hippocampus. **CA1.** a) ΔF/F time series and time-average color-coded room frame and arena frame-specific activity heat maps of a 13-unit subset from an 81-neuron ensemble recorded during pretraining on a familiar rotating arena. Activity localized in the arena frame can be dispersed in the roomframe (cell 1), and vice versa (cell 6), but typically, activity is dispersed in both frames. b) Ensemble spatial-frame-specific momentary positional information (I_pos_) is computed at 133 ms resolution in the room and arena frames. The time series of the difference (ΔI_pos_) indicates alternating preferences to signal information about the position in the room (positive ΔI_pos_) and in the arena (negative ΔI_pos_). During this 3-min episode, that information fluctuates between the two spatial frames approximately equally. c) The corresponding ensemble-summed firing rate time series. d) Overall spatial frame ensemble preference (SFEP) of 5 mice is quantified by the relative proportion of time the ensemble activity is room- or arena-preferring. While there is no preference during pretraining, CCT causes a preference for signaling arena-frame locations to emerge during the initial trial. The overall arena-preference maintains for at least a week and is expressed during the memory retention test. e) The corresponding spatial distribution of the SFEP in the room-frame shows that during CCT, discharge near the shock zone is roompreferring, whereas it is arena-preferring far from the shock zone, demonstrating goal-directed cognitive control in the spatial frame preference of hippocampal discharge.

We then investigate whether such cognitive control training causes learning to learn by comparing performance during initial training (Ti) and subsequent training (Ts) in a novel environment (Fig. 2a). Mice were initially trained to perform one of three task variants on a rotating arena (Fig. 2S1). The cognitive control training (CCT) group was trained in the “R+A-“ task variant, so named because use of room-frame spatial cues is reinforced (R+) while unreinforced arena-frame spatial cues should be ignored (A-) ^23^. Mice in the place learning (PL) control group were trained in the “R+” task variant that differs from the R+A-task conditions only by the presence of shallow water on the arena surface that attenuates arena-frame olfactory cues, sufficient for recognizing places on the arena ^23,25^. Accordingly, R+ training requires less cognitive control to ignore the task-irrelevant arena locations of shock. Shock was never turned on for the spatial exploration (SE) group that otherwise experienced the identical physical conditions as the CCT mice. After initial training, all mice were subsequently trained in a novel environment to do the same R+A-task that requires cognitive control (Fig. 2a).

**Figure 2:**
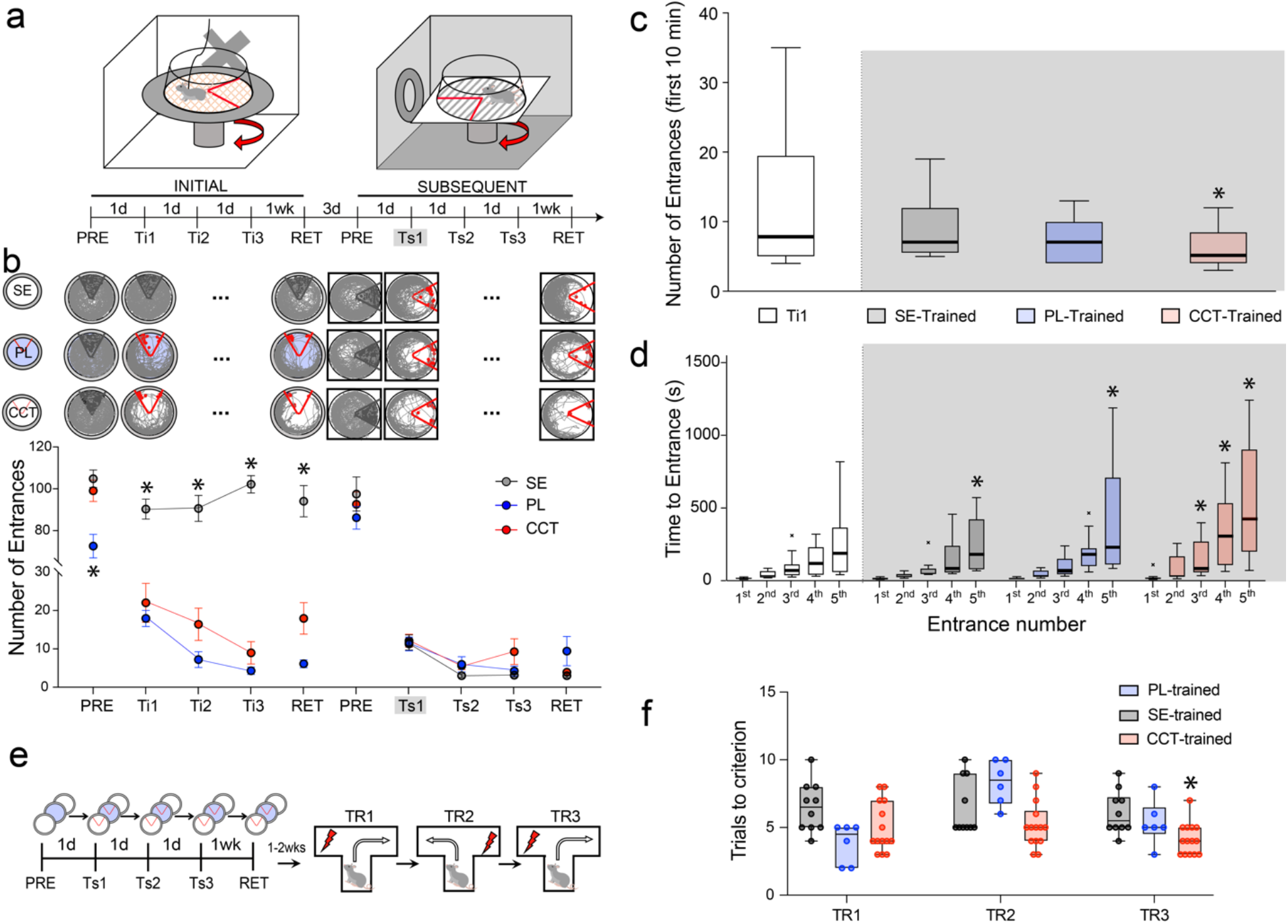
Learning to Learn. a) Behavioral protocol and schematic of active place avoidance in the distinct INITIAL and SUBSEQUENT training environments. After pretraining (PRE, shock off, indicated by black), mice received three daily training trials with shock on (indicated by red) in the initial (Ti) and subsequent (Ts) environments. A week later, each mouse received a retraining (RET) trial to evaluate 1-week memory. The environments were counterbalanced, and each trial was 30 min. There were 3 groups of mice: CCT (cognitive control training n = 13), PL controls (place learning n = 11), and SE (spatial exploration, untrained controls n = 9). During Initial trials, the CCT and PL controls received shock in the shock zone; SE mice were not shocked. For PL mice, shallow water on the arena floor attenuates the salience of task-irrelevant olfactory cues. In the subsequent training environment, all mice received the same R+A-place avoidance training in the same conditions. b) Example paths during select trials of the same representative mice; dots indicate the mouse’s location when it was shocked (red), and when it would have been shocked if the shock was on (black). The group average performance measured during 30-min by entrances into the shock zone or equivalent area for the SE mice. Statistical analysis of subsequent training (see Supplemental Information for analysis of initial training): Group X Trials1-3 two-way RM ANOVA, Group: F_2,30_ = 0.02; p = 0.99; Trials: F_2,56.7_ = 13.5, p = 10^−5^, η^2^ = 0.19, Tukey-HSD: Ts1 > Ts2 = Ts3; Interaction; F_4,65.2_ = 1.15, p = 0.34). The groups differed on the 1-week retention test (F_2,29_ = 3.77, p = 0.04, η^2^ = 0.21). *p < 0.05 comparing the corresponding group measures. c,d) CCT performance during the first trial in the initial environment (Ti1) compared to performance during the first trial in the subsequent environment (Ts1). *p < 0.05 compared to the SE group, and the SE group’s initial entrance times. c) The number of shock zone entrances during the first 10-min epoch. CCT Ti1 vs Ts1: paired *t_12_* = 2.23, p = 0.02, d = 0.62. CCT Ti1 vs PL Ts1: *t*_22_ = 1.56, p = 0.13; CCT Ti1 vs SE: *t*_20_ = 1.02, p = 0.32) d) The times of the first 5 shock zone entrances during the R+A-tasks. Group X Entrance1-5 two-way ANOVA, Group: F_2,62.9_ = 6.26, p = 10^−3^, η^2^ = 0.06; Entrance Number: F_4,52.7_ = 34.2, p = 10^−11^, η^2^ = 0.40; Interaction: F_8,66.4_ = 1.88, p = 0.07; Tukey-HSD tests confirmed that compared to the time of the first entrance, the CCT group had significantly increased the time to enter the shock zone by their third entrance, whereas the other two groups had only done so by their fifth entrance. CCT Phase (Ti1, Ts1) X Entrance Number two-way RM ANOVA, Phase: F_1,50_ = 14.6, p = 10^−4^, η^2^ = 0.10; Entrance Number: F_4,50_ = 9.81, p = 10^−3^, η^2^ = 0.31; Interaction: F_4,50_ = 2.41, p = 0.06) e) Learning to learn during initial CCT generalizes to L/R discrimination in a T-maze. T-maze alternation in 10-trial blocks was trained 7 to 14 days after either CCT (n = 14), PL (n = 6), or SE training (n = 10). f) Performance measured as the trials to reach the criterion of the second correct responses without receiving any shock. Group X Trial two-way RM ANOVA Group: F_2,27_ = 12.4, p = 10^−3^, η^2^ = 0.48; Trial: F_1.902,51.36_ = 1.86 p = 0.17; Interaction: F_4,54_ = 1.42, p = 0.24, Sidak’s multiple comparisons of session 3 performance: CCT < SE = PL). *p < 0.05 compared to all others.

## Subsequent Learning After Initial Training

After characterizing the three groups and their differences during pretraining and initial learning (Fig. 2a,b; see Supplemental Information subsection “Pretraining and Initial Learning”), we compared them during subsequent training in a novel environment when the conditions were identical for all groups. During pretraining, the mice did not express the thigmotaxis or avoidance from initial training, demonstrating the mice treat the environment as novel (Fig. 2b). During subsequent training, the groups did not differ in how much they walked (Fig. 2S2a); all mice learned the new R+A-place avoidance rapidly. When 30-min trials were assessed, the groups could not be distinguished by errors (Fig. 2b), or by the time to first enter the shock zone (Fig. 2S2b), although the CCT group tended to make fewer errors on the retention test 1 week later (Fig. 2b).

Because learning was rapid and may have been effectively complete within the first 30 min, we examined avoidance during the first 10-min of the first subsequent training trial (Ts1). The CCT mice entered the shock zone less during subsequent (Ts1) compared to initial training (Ti1; Fig. 2c). During Ts1, the PL and SE groups performed indistinguishably from the CCT group during Ti1 (Fig. 2c), suggesting that unlike the CCT group, their initial experiences did not alter subsequent learning. The times at which the mice entered the shock zone during Ts1 revealed that CCT mice learned to avoid entering the shock zone better than the other groups (Fig. 2c). The CCT mice also learned the subsequent avoidance faster than they learned the initial avoidance, as assessed by comparing the times of the first 5 entrances during Ti1 and Ts1 (Fig. 2d, Fig. 2S3a). Analysis of the number of entrances during the six 5-min epochs of Ti1 and Ts1, the initial and subsequent first training sessions, further confirms that CCT mice learn to learn (Fig. 2S4). The conclusion that the CCT group learned to learn during initial training is further supported by a significant correlation between the strength of place avoidance at the end of initial training (trial RET) and the strength of place avoidance at the start of subsequent training in the entirely novel environment, which was not observed for the PL group (Fig. 2S5). Indeed, the correlations between conditioned avoidance during the initial trials (Ti1-3) and during subsequent (Ts1) training increased from Ts1 to Ts3, but not within the PL group (Fig. 2S5c).

## Learning to Learn

CCT also improved subsequent learning of a left-right (L/R) discrimination task in a novel T-maze environment (Fig. 2e), the reversal of which is sensitive to hippocampus function ^26^. Mice discriminated equally well during the first session of 10 trials whether they had received initial CCT, PL or SE training at least a week before, but performance of the CCT mice tended to be better when the reinforcement was reversed for session 2 and was significantly better when it was reversed again in session 3 (Fig. 2f). Unlike the first session, the reversals require mice to distinguish between recalling the prior and current safe versus punished arms. These data together demonstrate that learning to learn in the CCT group persists for weeks and generalizes beyond the place avoidance task, whereas place learning itself in the PL group does not.

## CCT persistently changes neural circuit function

We confirmed that the dentate gyrus (DG) is crucial for expressing active place avoidance following CCT by targeting bilateral muscimol inactivation to DG (Fig. 3S1). Accordingly, to assess whether CCT training modifies entorhinal-hippocampus circuit function, a subset of the mice had been implanted with stimulating electrodes in the medial perforant path (MPP) input to DG and CA1 along with silicon 32-site linear electrode arrays that spanned the somatodendritic axis of dorsal hippocampus (Fig. 3S2,3). We confirmed electrode targeting of the MPP using a transgenic mouse that predominantly expresses the inhibitory DREADD hM4Di in layer 2 of the medial entorhinal cortex (MEC) under control of the tTA-TetO system to attenuate MPP stimulation ^27^ (Fig 3S3).

We evaluated whether the CCT, PL, and SE experiences change MEC-DG circuit function by measuring source-localized DG evoked potential responses to 0-250 μA MPP test stimulation before pretraining and before the 1-week memory retention session, (Fig. 3a, 3S4). PL (n=5) and SE (n=5) did not cause detectable changes, but CCT (n=7) reduced the source-localized field excitatory postsynaptic potential (fEPSP) slope in the molecular layer of the suprapyramidal division of DG (supraDG; Fig. 3b). Changes were minimal in the population spike and at the infrapyramidal division (infraDG, Fig. 3c). Electrode impedances, the current source density (CSD), and theta power did not change systematically (Fig. 3S5). CSD analysis shows that CCT reduces the current sink in the response at the middle molecular layer of supraDG, where the MPP terminates (see Fig. 4c) ^28^. These changes occur 2 hours after the first CCT session and plateau after the second trial (Fig. 3S6). These CCT-induced functional changes persisted at least 60 days without further training (Fig. S37).

**Figure 3:**
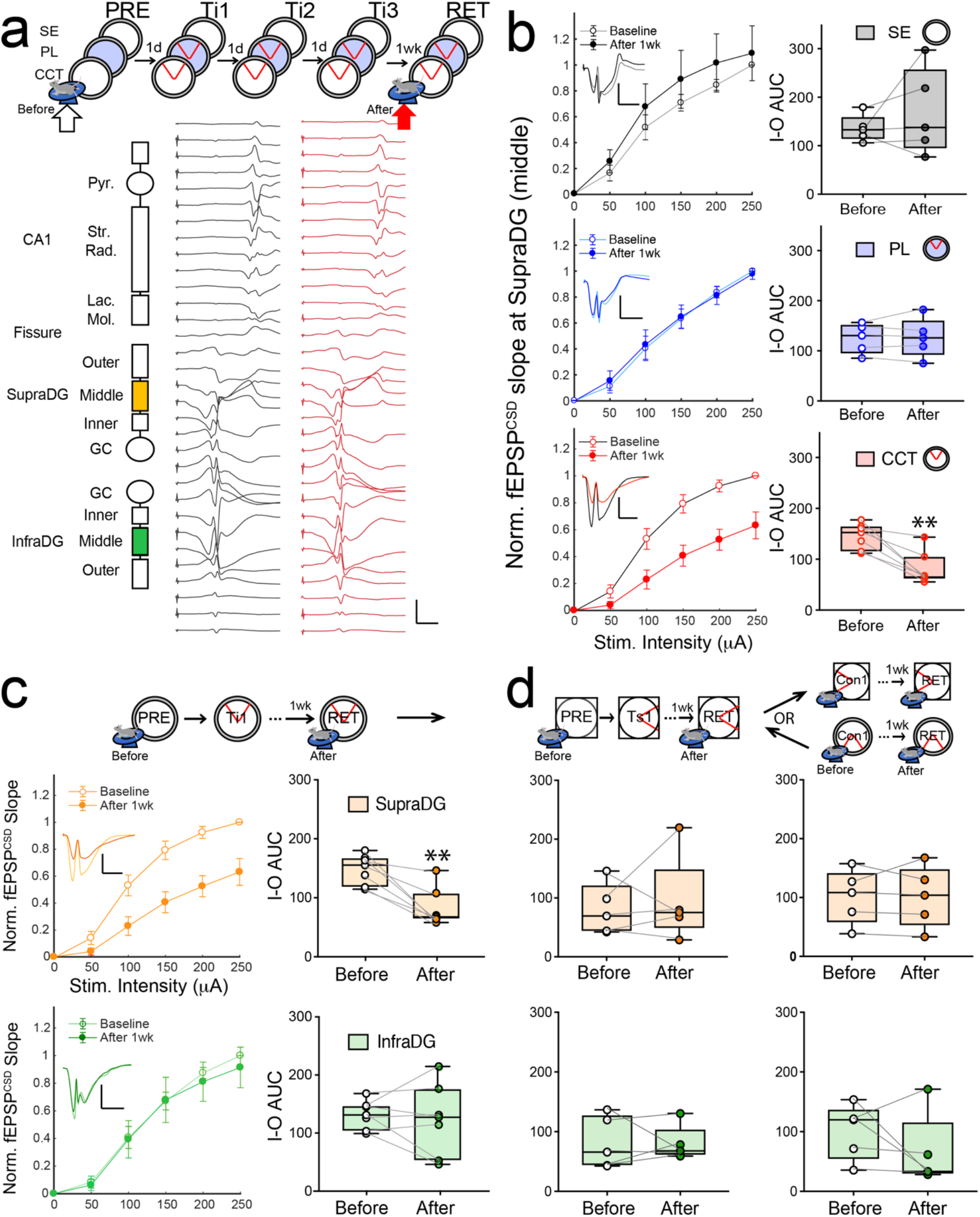
Cognitive control training persistently changes entorhinal-dentate circuit function. a) Experimental design and example of a raw voltage trace evoked response recorded on the 32-ch linear silicon probe. Schematic depicts that mice receive one of three types of training in the same environment and before each training session, evoked responses to perforant path stimulation are recorded while the mouse runs on a wheel. The voltage sources are localized to synaptic compartments along the somato-dendritic axis using current source density analyses. Scale bar: 200 μA/mm^3^, 5 ms). b) Input-output curves showing that that CCT (n = 7) but not PL (n = 5) or SE (n = 5) training attenuates the supraDG population synaptic response to MPP stimulation in the middle molecular layer (Group X Training RM ANOVA, Group: F_2,14_ = 1.64, p = 0.23; Training: F_1,14_ = 0.82, p = 0.38; Interaction: F_2,14_ = 6.90, p = 0.008, η^2^ = 0.50; Sidak’s multiple comparisons: only CCT changes circuit function **p = 0.007). (scale bar: 50 μA/mm^3^, 5 ms) c) CCT caused changes at the corresponding middle molecular layers of the supraDG (repeated from b left) but not the infraDG response to MPP stimulation (Area X Training RM ANOVA, area: F_1,12_ = 0.49, p = 0.50; training: F_1,12_ = 11.08, p = 0.006, η^2^ = 0.48; interaction: F_1,12_ = 6.69, p = 0.02, η^2^ = 0.36; Sidak’s multiple comparisons: only SupraDG changes circuit function **p = 0.003; supraDG was significant paired t_6_ = 5.09, **p = 0.002, d = 1.92) but not at infraDG sites (paired t_6_ = 0.46, p = 0.66). Scale bar: 50 μA/mm^3^, 5 ms. d) No further changes are observed in the synaptic responses after either additional CCT in a novel environment (supraDG: paired t_4_ = 0.45, p = 0.67; infraDG paired t_4_ = 0.12, p = 0.90) or in the same environment with changed shock zone (supraDG paired t_4_ = 0.02, p = 0.98; infraDG paired t_4_ = 1.13, p = 0.32).

**Figure 4:**
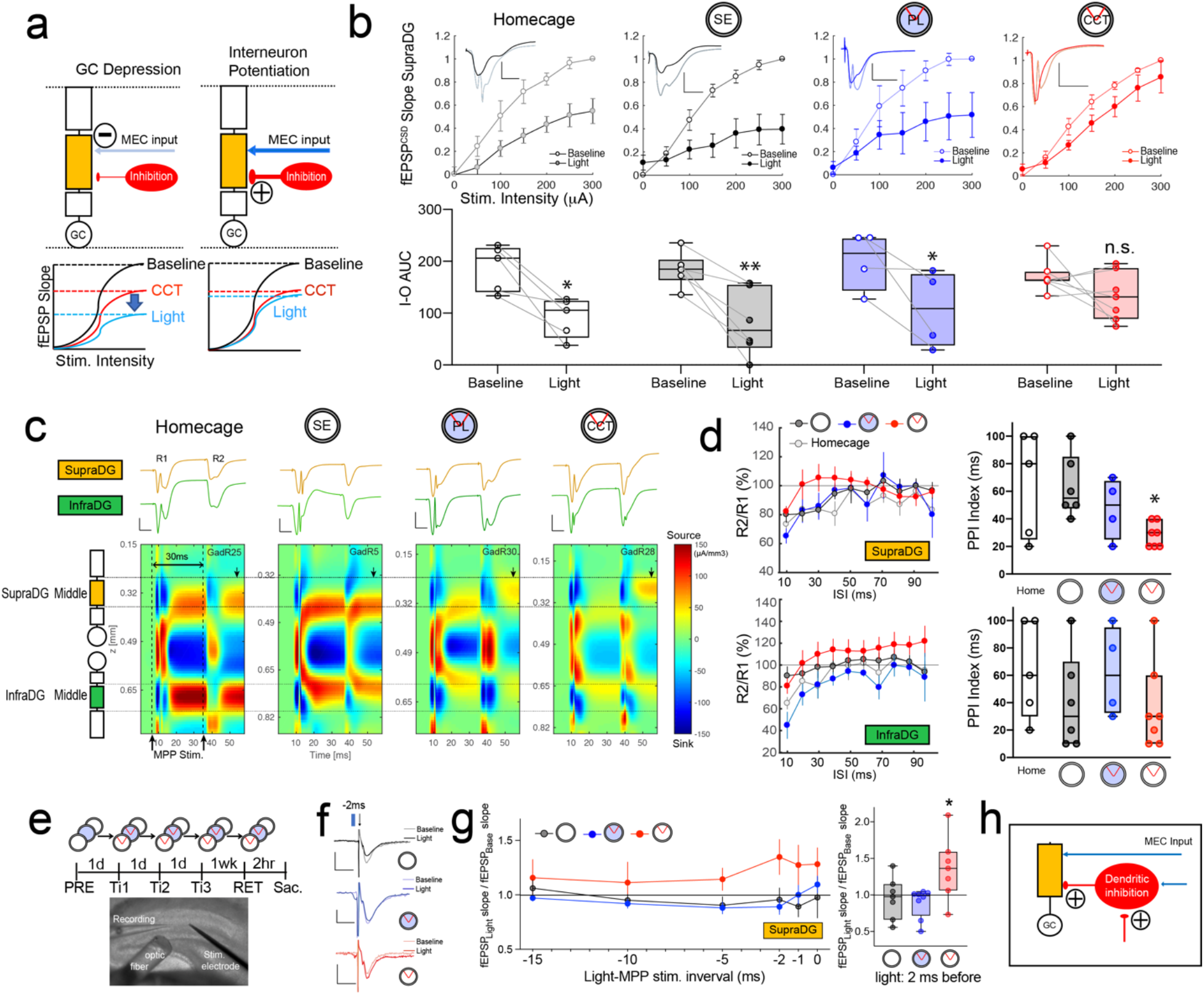
Cognitive control training persistently changes inhibitory entorhinal-dentate circuit function. a) Schematic of MPP→granule cell (GC) depression and potentiation of GC inhibition, two hypotheses to explain the reduced DG response to MPP stimulation that is caused by CCT. b) One-week post-training, input-output curves (above) and their summary as the area-under-the curve (below) quantify the fEPSP at supraDG in response to MPP stimulation under urethane anesthesia, with and without concurrent light activation of ChR2 expressed in Gad2+ cells. Light-induced inhibition decreased responses, which mimics the effect of training in home cage (n = 5), SE-trained (n = 6) and PL-trained (n = 4) control mice but the effect is occluded in CCT-trained mice (n = 7) (Group X Light stimulation RM ANOVA, Group: F_3,18_ = 0.40, p = 0.75; Light stimulation: F_1,18_ = 36.50, p < 0.0001, η^2^ = 0.67; Interaction: F_3,18_ = 1.46, p = 0.26; Sidak’s multiple comparisons: Homecage: p = 0.01, SE: p = 0.003, PL: p = 0.03, only CCT occluded the light stimulation effect. p = 0.75). Scale bars: 100 μA/mm^3^, 10ms. c) Current-source density examples of single responses to paired-pulse inhibition in homecage and SE-trained control mice and CCT mice (Scale bars: 100 μA/mm^3^, 5 ms), illustrate that MPP-stimulated inhibition is attenuated in the termination zone of supraDG after CCT. d) Summary of the training group PPI curves and indices (supraDG: F_3,18_ = 3.22, *p = 0.04, η^2^ = 0.15; infraDG: F_3,18_ = 0.95, p = 0.43). e) Experimental design and photomicrograph of *ex vivo* hippocampus slices prepared 1 week after training, showing the optic fiber and recording and stimulating electrodes. fEPSP responses to stimulation of MPP terminal fibers were recorded with 1-ms light pulses to activate ChR2 in Gad2+ cells. f) Exemplary responses from SE-trained mice (black), PL-trained mice (blue) and CCT-trained mice (red) with and without (baseline) light-activation of ChR2 in Gad2+ cells 2-ms pre-stimulation (below). Scale bars: 10 mV, 5 ms. g left) Summary of the ratio of the fEPSP slope response measured with light stimulation normalized to the baseline response at supraDG (Group X Light-Stim ISI RM ANOVA, Group: F_2,19_ = 2.67, p = 0.09; Light-Stim ISI: F_5,95_ = 1.43, p = 0.22; Interaction: F_10,95_ = 0.95, p = 0.49). g right) Light activation 2 ms before MPP stimulation caused increased supraDG synaptic population responses in CCT-trained mice (n = 7) compared to SE-trained mice (n = 7) or PL-trained mice (n = 8) (ANOVA F_2,19_ = 4.48, p = 0.025, η^2^ = 0.48). h) Schematic of hypothesized mechanism of MPP circuit changes resulting in MPP response suppression, increased feedback inhibition, and increased disinhibition. Averages of mice are presented, 1-2 slices per mouse were studied.

We verified that these CCT-induced changes are not the direct result of the MPP stimulation by exploiting naturally occurring, highly-synchronous MPP-originating activations known as type 2 dentate spikes, or hereby referred to as DS_M_. We reasoned that CCT should change DS_M_ events like we observed from experimental MPP stimulation. As predicted, the slope of the MPP-associated current sink components of DS_M_ decreased in supraDG but not in infraDG (Fig. 3S8).

If the changes in neural circuit function caused by CCT are due to acquiring a content-specific place avoidance memory, then learning an additional place avoidance should cause additional changes, whereas if the neural circuit changes result from the “process memory” of learning to learn ^29^ then the additional training should not cause incremental neural circuit changes. A subset of initial CCT mice received subsequent CCT to acquire a different place avoidance memory in the same environment with a different shock location (conflict, n = 5) and/or in a novel environment (n = 5). The 4 mice/group that received both types of subsequent training, received conflict or novel environment training in counterbalanced order. Although subsequent learning was better than initial learning (Fig. 3S9), the subsequent training experiences did not cause incremental changes in neural circuit function (Fig. 3d), consistent with a change in process rather than content-specific memory.

## CCT changes inhibitory circuit function

We hypothesized that CCT changes excitation-inhibition coordination at supraDG. The conjecture was based on the following: 1) DG interneurons strongly regulate principal cell firing ^30,31^. 2) CCT increases the sub-second coupling between action potential discharge of excitatory granule cells and inhibitory neurons in DG ^21^. 3), and spatial learning increases DG mossy fiber-mediated feedforward inhibition ^32^. Indeed, the dentate gyrus reveal more inhibitory neurons in supraDG than infraDG in Gad2Cre-ChR2-eYFP mice that express eYFP in GABAergic neurons (Fig. 4S1) as has been reported ^33,34^. The Gad2Cre-ChR2-eYFP mice were used to distinguish two hypotheses for the weakened supraDG response to MPP stimulation (Fig. 4a). Under urethane anesthesia that reduces ongoing activity ^35,36^ (Fig. 4S2), channelrhodopsin activation of Gad2-expressing DG cells in the vicinity of the recording site, blocked population spike responses in all mice when light stimulation preceded MPP stimulation by 5 ms, indicating effectiveness of the inhibition and attenuation of potential feedback responses (Fig. 4S3). Importantly, activating this inhibition 5 ms before MPP stimulation (but not at the same time; Fig. 4S4), mimicked the CCT-induced attenuation of MPP responses at supraDG of task-naïve mice, and a week after the SE or PL experiences. In contrast, the inhibition-mediated attenuation was occluded at supraDG of the CCT mice, consistent with the hypothesis that CCT potentiates inhibition in supraDG (Fig. 4b). Current Source Density (CSD) analysis of the response to paired-pulse stimulation reveals a delayed source at the supraDG MPP terminals after the second pulse. This delayed source is especially strong in CCT mice (Fig. 4c), suggesting that CCT persistently enhances activity-dependent inhibition at the MPP termination. Indeed, compared to task naïve mice, CCT mice, but not SE and PL mice, significantly decrease MPP-stimulated interneuron-mediated inhibition at supraDG (Fig. 4c,d), but not at infraDG or at the granule cell layer (Fig. 4S5). These findings indicate that CCT alters pathway-specific excitation-dependent inhibition at supraDG not only by generally increasing inhibition, but perhaps also by causing disinhibition at the MPP terminals.

To test the possibility that CCT disinhibits MPP inputs, we exploited the strong reduction of spontaneous activity in *ex vivo* hippocampus slices, and used slices from Gad2Cre-ChR2-eYFP mice to directly stimulate inhibition in the Gad2-expressing cells within supraDG (Fig. 4e). Concurrent light activation of Gad2-expressing cells inhibits the response to MPP stimulation (Fig. 4S3), whereas the activation 2 ms before MPP stimulation increases supraDG synaptic population responses in slices from CCT mice but not at infraDG, nor in slices from PL or SE mice. This demonstrates local disinhibition of MPP inputs to supraDG after CCT (Fig. 4f,g, 4S6). We also observed that the fEPSP responses to MPP stimulation in *ex vivo* slices significantly strengthen at supraDG after PL but not after SE or CCT (Fig. 4S7). Together these findings indicate that while CCT globally increases inhibition of supraDG in the freely-behaving mouse, CCT also disinhibits strong MPP inputs to supraDG.

## An entorhinal-hippocampal subcircuit tuned by cognitive control training

We find that CCT (Fig. 1) results in learning to learn that generalizes to other tasks in which the explicit information content is novel (Fig. 2), and that CCT causes pathway-specific suppression and enhancement of the synaptic response to MPP activation (Figs. 3, 3S7, 4). These particular subcircuit changes can mediate an input-controlled excitation-inhibition coordination for increasing the signal-to-noise ratio by globally potentiating inhibition while disinhibiting the currently strongest inputs. This neural network motif will also stabilize the connection’s inputoutput relationship ^37,38^. Similar disinhibitory plasticity has also been described at the Schaffer collateral and other cortical synapses ^39–41^, where CCT causes persistent increased expression of PKMζ, which is itself necessary and sufficient for maintaining LTP ^17,42^. The CCT-induced subcircuit changes were detected in field potential recordings, indicating they are not the sparse changes that mediate content-specific memory formation that the synaptic plasticity and memory hypothesis predicts ^43^. Indeed, the functional changes we identified are more consistent with the related concepts of learning to learn, learning set and memory schema, and suggest a mechanism for the increased sub-second coupling of place cell and inhibitory cell discharge that was reported in dentate gyrus during moments of memory discrimination ^21^. Together these findings suggest a role of dentate gyrus in judiciously and flexibly representing environmental features to which internally-generated spatial representations are transiently registered, distinct from a role in pattern separation ^44–46^. Furthermore, the present findings motivate investigating whether learning to learn causes similar excitation-inhibition changes at other hippocampal and cortical sites. As such, our findings challenge interpretations of studies that are designed to investigate the neurobiological changes that underlie content-specific memory, because as we demonstrate, neurobiological changes that mediate generalized process memory impacting information processing can overlap with physiological changes that mediate the content-specific memory information storage itself.

The CCT-induced, persistent changes in entorhinal-hippocampus circuit function demonstrate that appropriately structured cognitive experiences can persistently change information processing within neural subcircuits. As such, the present findings strongly support the neuroplasticity hypothesis, although of course, CCT causes additional changes ^14,17,47^, with others yet to be discovered. Prior work showed that preemptive CCT in adolescence prevented cognitive impairment due to neonatal neurodevelopmental brain damage ^48^, and as also demonstrated here, mere place learning was not sufficient to cause several neurobiological changes that CCT causes. This highlights the importance of cognitive control training itself, and potentially how the practice of successfully ignoring salient distracting information can have widespread beneficial effects because of the additional cognitive challenge that CCT presents over navigation itself (Fig. 1), which has been exploited for effective CBT and learning to learn ^5,10^. These findings also demonstrate that common neurobiological mechanisms may underlie the traditionally separate fields of studying schema in cognitive psychology, CBT in psychotherapy, and learned information processing in neurobiology.

## Acknowledgements

Supported by NIH grants R01MH115304, R01NS105472, and R01AG043688.

## SUPPLEMENTARY INFORMATION

### Experimental Design - Environments

**Figure 2S1.**
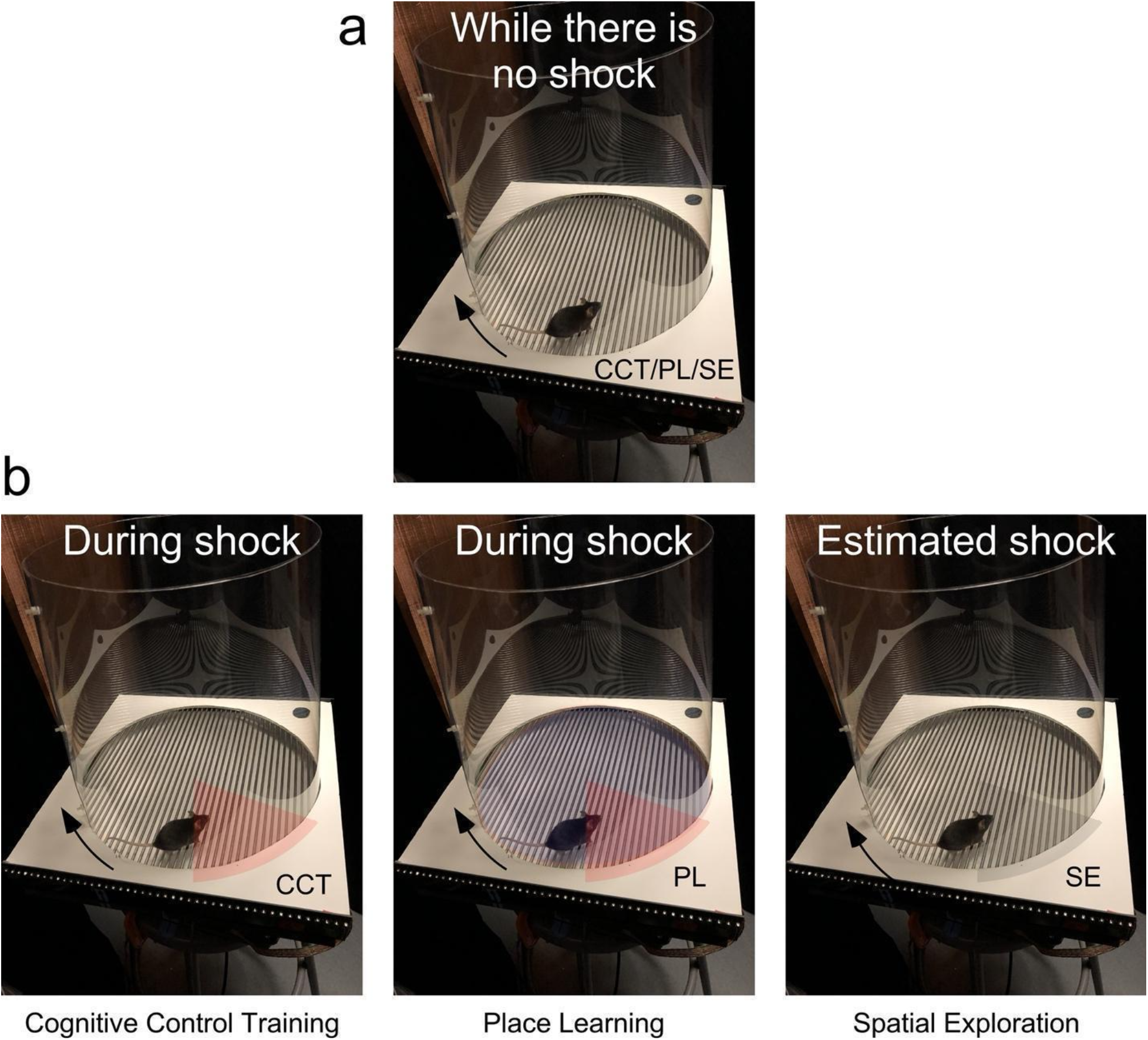
(related to Figure 2). Task variants. a) The apparatus and environment were essentially the same for the three task variants. More than 99% of the time in the apparatus, when there was no shock, the visual environment was identical across the tasks; only the place learning environment was different in that the arena surface was covered in shallow water. b) During the 500-ms shock the cognitive control training and place learning groups experienced an unpleasant foot shock (depicted as shaded sectors), whereas the spatial exploration group did not.

### Supplemental Results - Pretraining and Initial Learning Behavior

The CCT (n=13), PL (n=11), and SE (N=9) groups differ in the initial but not the subsequent pretraining trials with no shock, because mice walk less in the shallow water (Group x Trial twoway RM ANOVA; group: F_2,27_ = 27.7, p = 10^−7^; η^2^ = 0.49; Trial: F_1,50.5_ = 2.44, p = 0.12; interaction: F_2,50.5_ = 5.88, p = 0.005, η^2^ = 0.19; PL during initial < all other group x trials measures). This resulted in group differences in the number of times the mice enter the location of the future shock zone (Group x Trial two-way RM ANOVA, Group: F_2,31_ = 10.51, p = 10^−4^, η^2^ = 0.22; Trial: F_1,60.7_ = 0.042, p = 0.8; Group X Trial: F_2,60.7_ = 2.93, p = 0.06; Tukey-HSD: CCT = SE > PL), as can be seen in Fig. 2b. Consequently, performance in the initial and subsequent training sessions is analyzed separately.

During initial training the conditioned mice walk less and restrict themselves to the arena periphery compared to the SE mice; the conditioned mice did not differ in the distance they walked (Fig.2S2; Group x Trial two-way RM ANOVA, Group: F_2,41.2_ = 11.51, p = 10^−4^, η^2^ = 0.14; Trial: F_2,28.5_ = 1.16, p = 0.33, η^2^ = 0.002; Group X Trial: F_4,32.4_ = 5.83, p = 0.001, η^2^ = 0.03; SE > CCT = PL). Place avoidance learning measured as decreasing entrances into the shock zone was robust with significant effects of Group and the Group X Trial interaction (Fig.2b Group: F_2,45.6_ = 72.3, p = 10^−11^, η^2^ = 0.23; Trial: F_2,34.7_ = 4.63, p = 0.02, η^2^ = 0.006; Interaction: F_4,39.3_ = 8.31, p = 10^−5^, η^2^ = 0.01; Tukey-HSD: SE > CCT = PL). The two conditioned groups showed similar learning (Fig 2S2, Group: F_1,59.2_ = 0.93, p = 0.34, η^2^ = 0.004; Trial: F_2,33.5_ = 39.5, p = 10^−9^, η^2^ = 0.20, Tukey-HSD Trial1 > Trial 2 > Trial 3; Interaction: F_2,33.5_ = 1.57, p = 0.2; η^2^ = 0.01). One-week memory retention was robust with an obvious effect of Group measured by entrances into the shock zone (see Fig. 2b; F_2,31_ = 101.3, p = 10^−14^, η^2^ = 0.73, Tukey-HSD: SE > CCT = PL) as well as the time to first enter the shock zone in the two conditioned groups with PL better than CCT (Fig 2S2b, Group: F_1,31_ = 3.25, p = 0.05, η^2^ = 0.21).

**Fig. 2S2.**
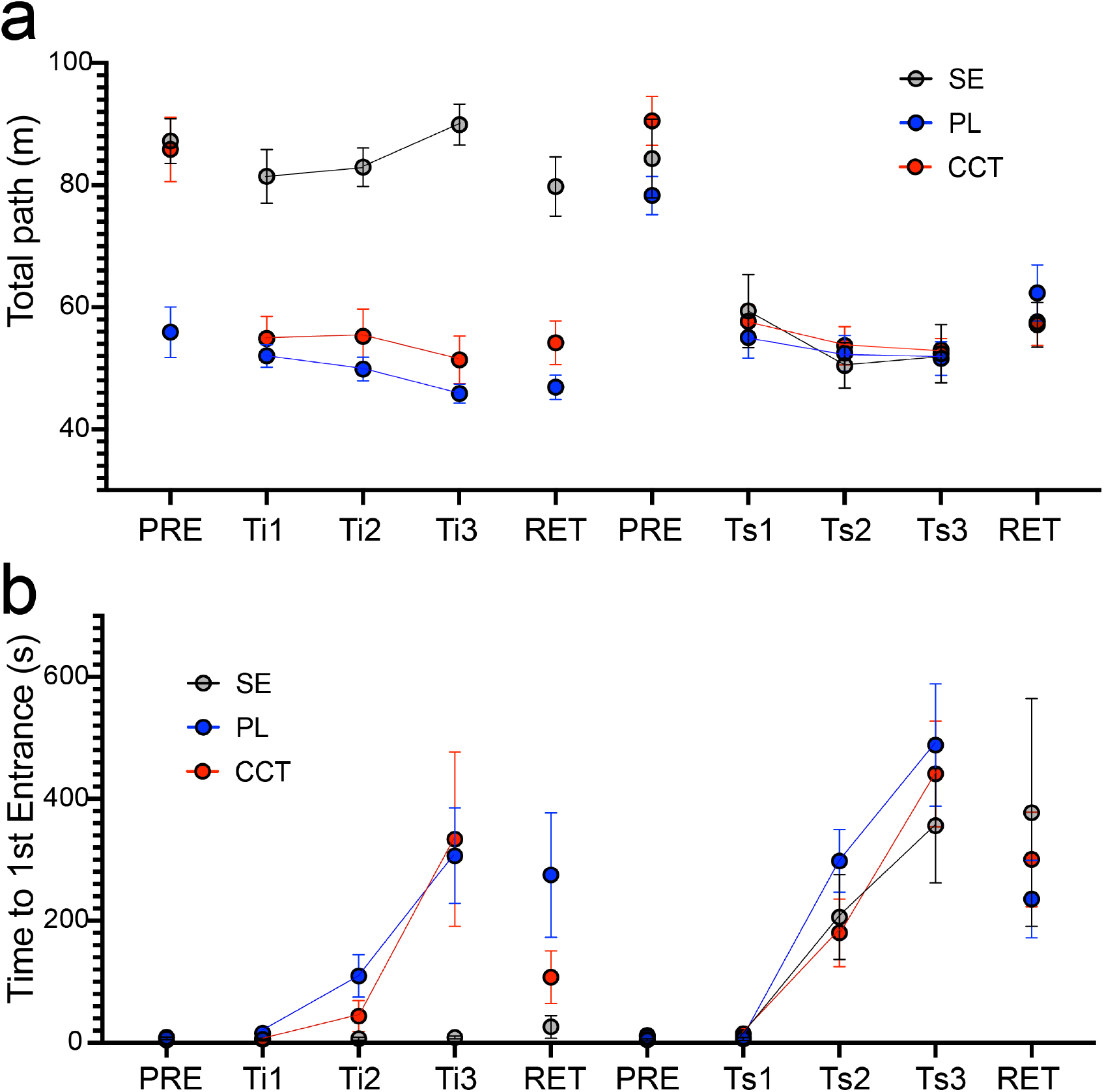
(related to Figure 2). Coarse measures of locomotion and place avoidance memory across the training protocol. **a)** Total path walked measures activity. b) Time to 1^st^ Entrance estimates between-session memory.

**Fig. 2S3.**
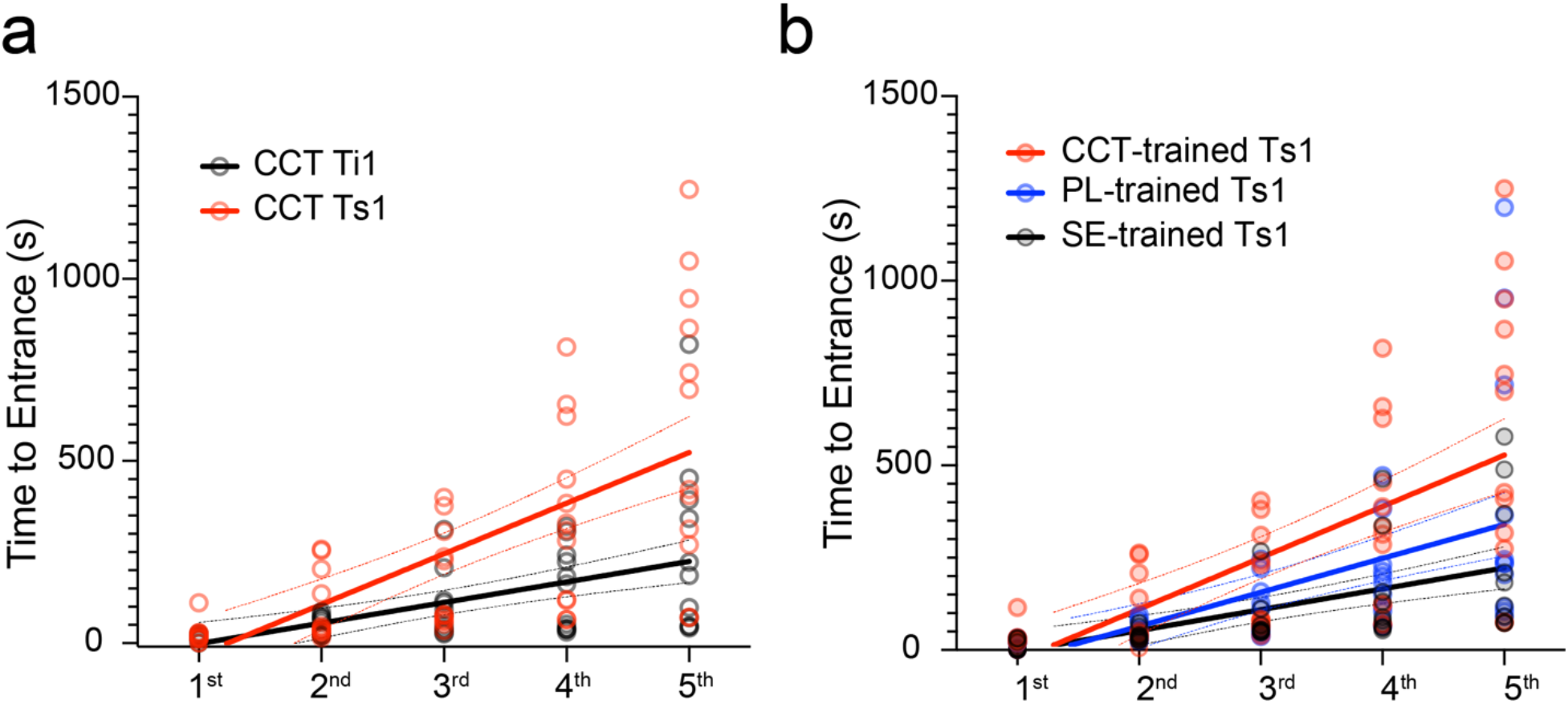
(related to Figure 2). Active place avoidance in a novel environment is improved by prior CCT training. a) Times to enter the shock zone for the first five times during subsequent training in entirely novel physical conditions (Ts1) are prolonged after initial training (Ti1) in the CCT group (comparison of slopes: F_1,111_ = 11.94, p = 0.0008, η^2^ = 0.10). b) CCT mice also prolonged entering the shock zone during subsequent training more than the other groups (comparison of slopes: F_2,159_ = 4.56, p = 0.01, η^2^ = 0.05). Solid lines depict linear regressions, and dotted lines depict 95% confidence bounds.

**Figure 2S4.**
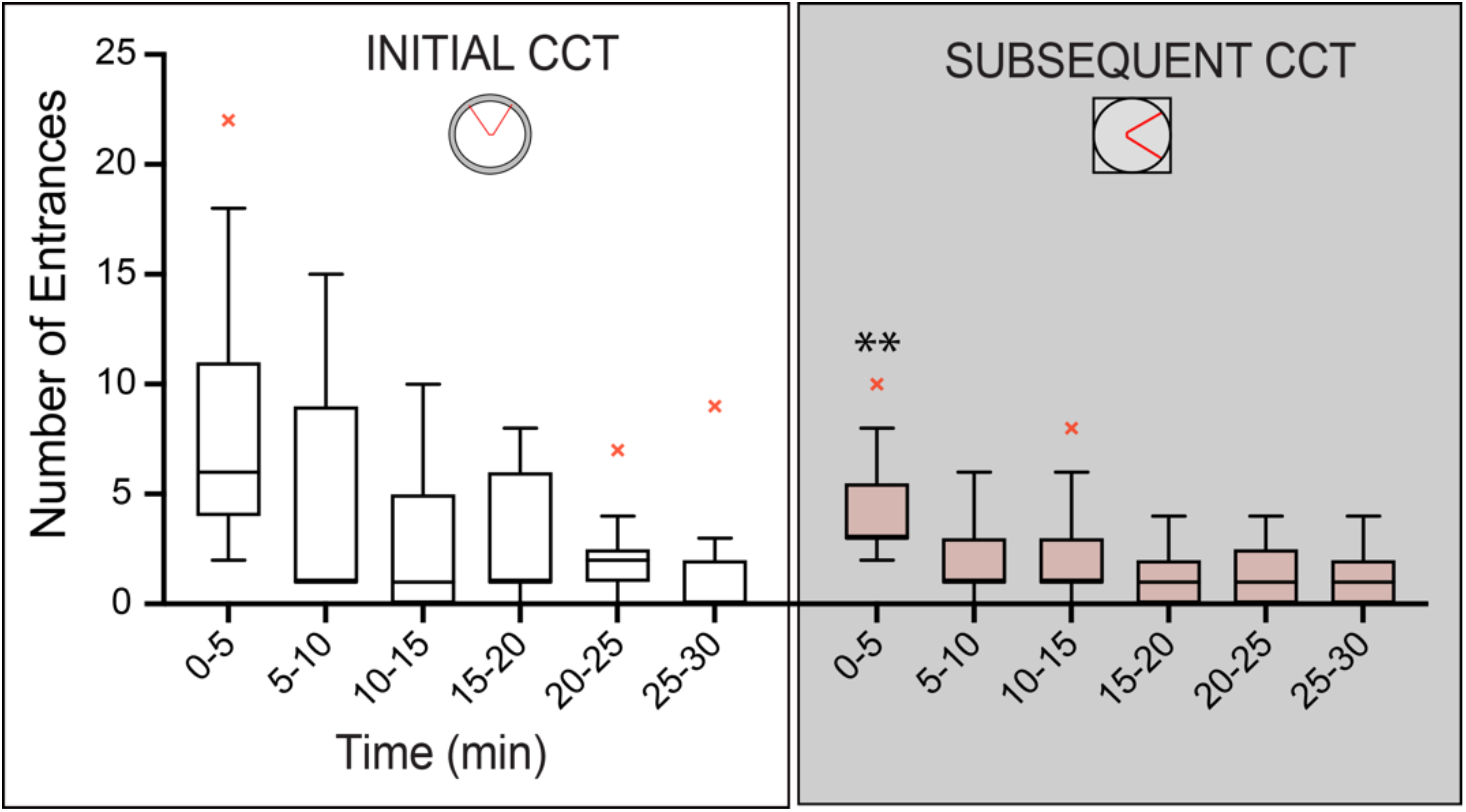
(related to Figure 2). Learning to learn. Within-session learning curve comparisons of the Initial and Subsequent CCT phases in different environments. Although there was no discriminative spatial information in common between the two environments, mice (n = 13) acquired the subsequent conditioned-place avoidance faster than the initial place avoidance, even though the sequence of experiencing the two environments was counterbalanced (Phase: F_1,72_ = 13.10, p = 0.005, η^2^ = 0.15; Time: F_5,72_ = 7.67, p < 10^−4^, η^2^ = 0.35; Phase X Time interaction: F_5,72_ = 1.53, p = 0.19, Sidak’s multiple comparisons 0-5min Initial CCT>Subsequent CCT **p=0.006). These data demonstrate learning to learn, as in the original description by Harlow (1949).

**Fig. 2S5.**
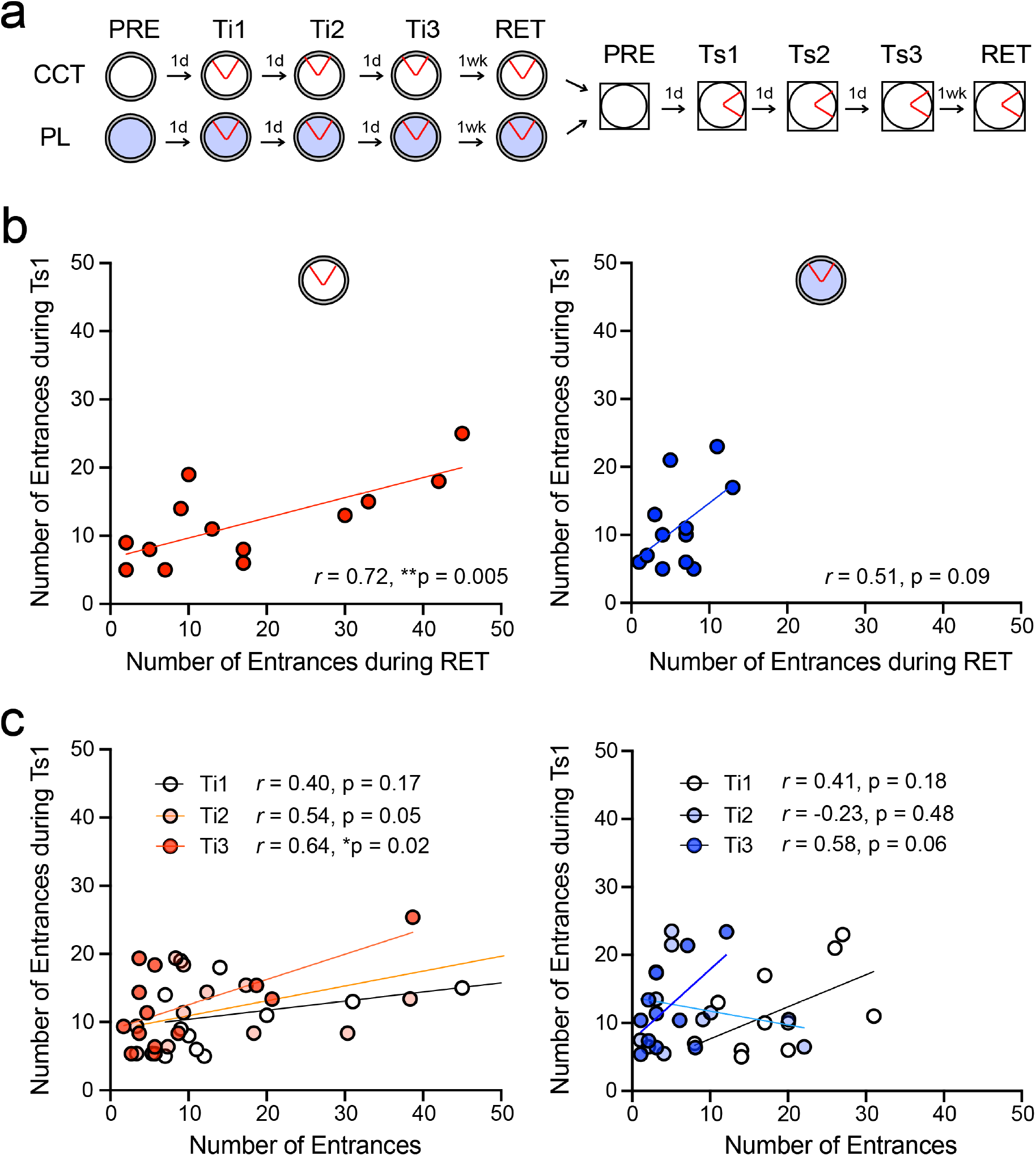
Learned place avoidance during initial training predicts subsequent place avoidance in a novel environment (related to Figure 2). a) Schematic training protocol. b) One-week retention of the conditioned place avoidance significantly predicts place avoidance in a novel environment amongst the CCT (left) but not the PL (right) mice. c) correlation between initial training trials (Ti1-3) and during subsequent (Ts1) training increased from Ts1 to Ts3 in CCT group, but not within the PL group.

### Supplemental Results - Technical and Experimental Validation

**Figure 3S1.**
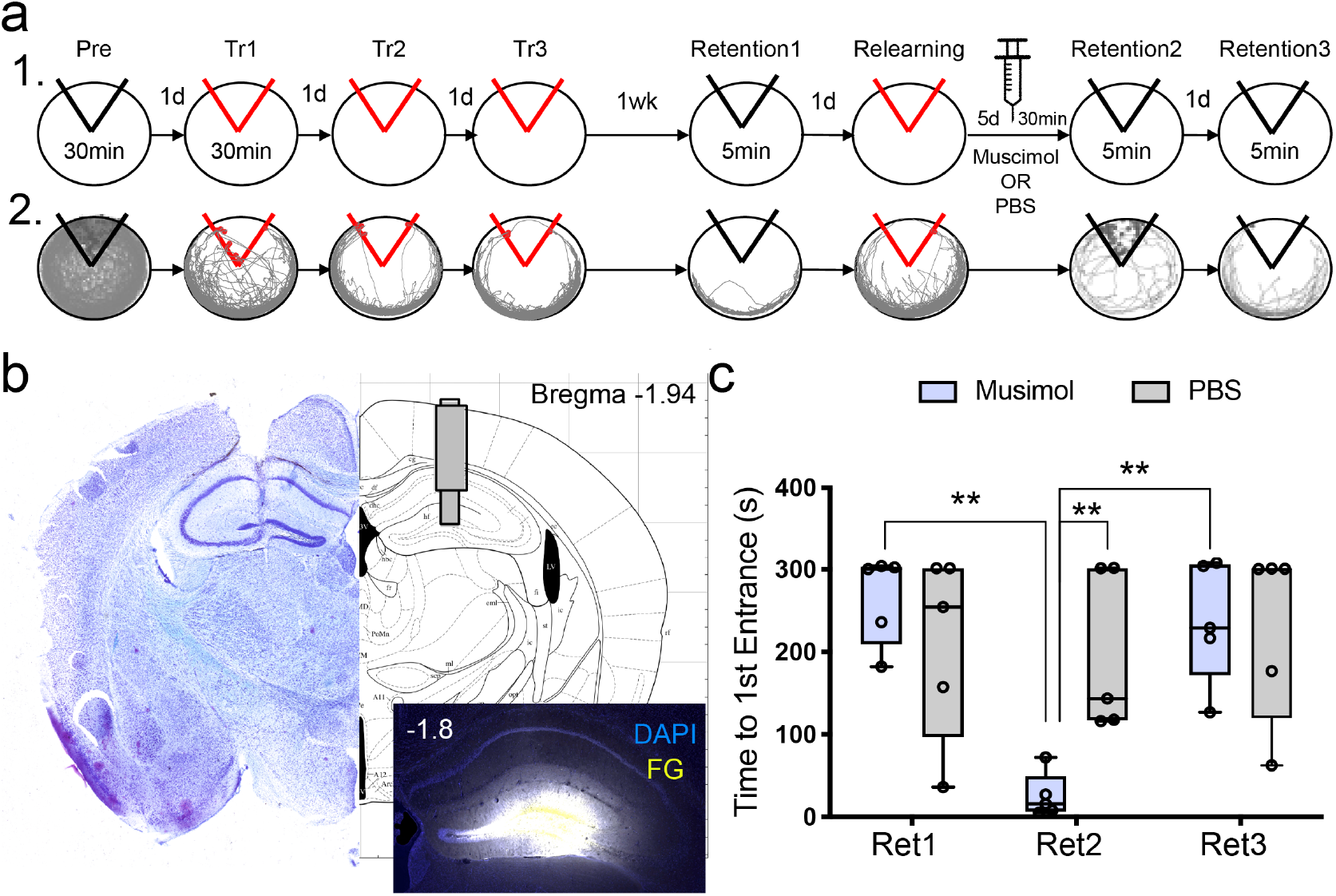
(related to Figure 3). Muscimol inactivation demonstrates DG is crucial for expression of active place avoidance several days after CCT training. a) Experimental design (1) and the tracks of an exemplar muscimol-injected mouse during each behavioral session (2). b) Schematic from Franklin and Paxinos^47^ illustrating the injection and guide cannulae target, a histology section of the cannulation track, as well as a histological section after injecting fluorogold (FG) to estimate the infusion spread. c) A measure of memory expression illustrates that targeting muscimol at DG reversibly impairs established active place avoidance memory (n = 5/group, Group X Retention session RM ANOVA Group: F_1,8_ = 2.46, p = 0.16; Retention session: F_2,16_ = 5.84, p = 0.01, η^2^ = 0.42; interaction: F_2,16_ = 4.01, p = 0.03, η^2^ = 0.33).

**Figure 3S2.**
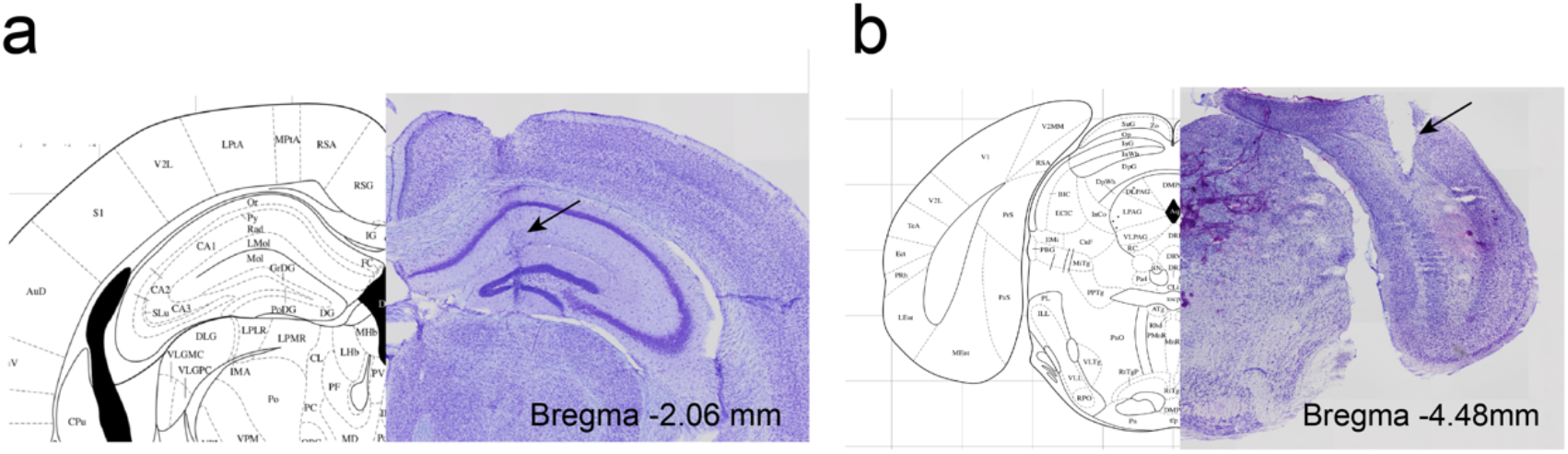
(related to Figure 3). Targeting of the recording and stimulating electrodes. Schematic and Nissl-stained histological sections from an exemplar mouse illustrating the a) recording and b) stimulation electrode sites. Schematics from Franklin and Paxinos^47^.

**Figure 3S3.**
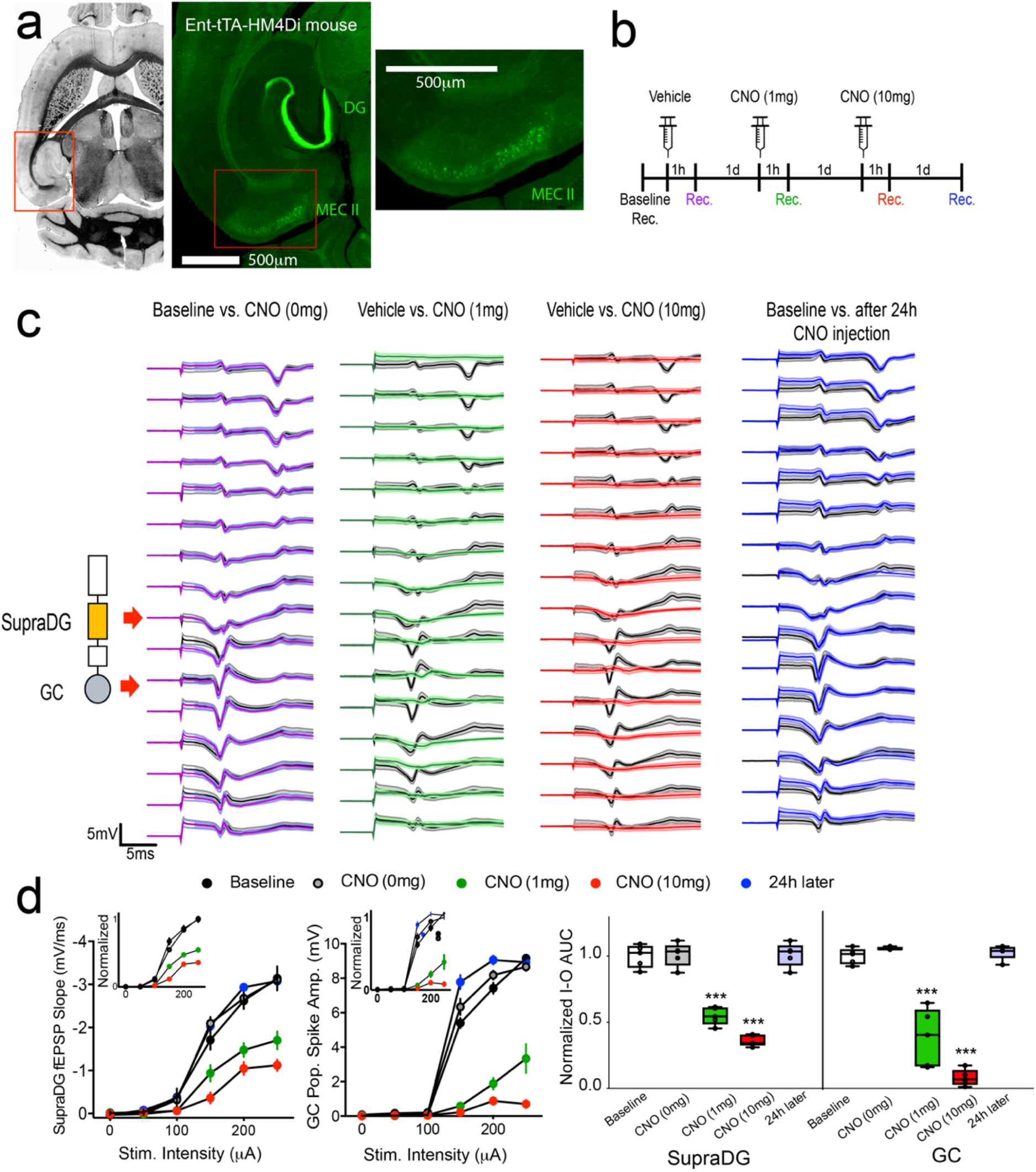
(related to Figure 3). Confirmation that electrical stimulation targeted the MPP. a) Nissl section and anti-human antigen (HA) immunostaining of transgenic mice (TRE-hM4Di-tetO+/- crossed with Ent-tTA+/-) that express the inhibitory DREADD under control of doxycycline withdrawal. The inhibitory DREADD immunoreactivity localizes to hippocampusprojecting MECII neurons. Immunofluorescence shows expression in MECII cell bodies, in the perforant path axons and in their termination zones in the molecular layer of the dentate gyrus and CA3. Expression is faint in CA1 *str. lacunosum moleculare*. Expression is in ~20% of MECII stellate cells and 5% in MECIII cells with less in adjacent regions (Kanter et al. 2017). b) Experimental design to evaluate effective stimulation of the MPP. c) Average evoked responses recorded with a 16-site linear silicon probe spanning the somato-dendritic axis of dorsal hippocampus. d) Input-output curves of fEPSP slope responses recorded from the DG molecular layer and population spike responses recorded from the DG hilus. These data quantify that responses to electrical stimulation targeted to the angular bundle is dose-dependently and reversibly suppressed by CNO-mediated activation of hM4Di in the MPP (Area X Dose RM ANOVA, Area: F_1,8_ = 7.55, p = 0.02, η^2^ = 0.49; Dose: F_4,32_ = 168.6 p = 10^−21^, η^2^ = 0.95; interaction: F_4,32_ = 5.78, p = 0.001, η^2^ = 0.42, Sidak-corrected within-area comparisons ***p<0.001).

**Figure 3S4.**
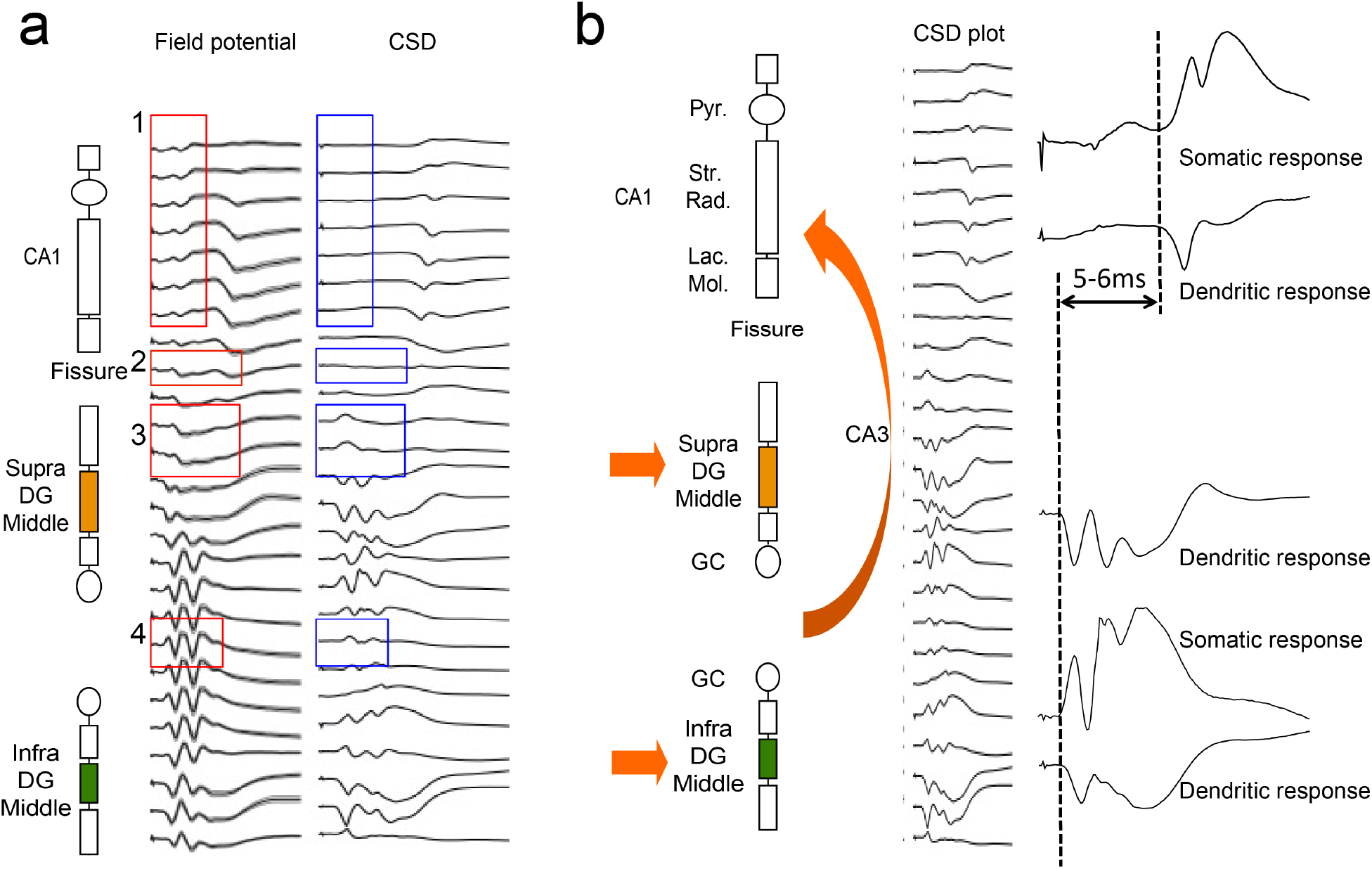
(related to Figure 3). Membrane potential vs. Current source density (CSD) plots to illustrate signal localization. a) comparison of the raw field potential (mV) plots and corresponding CSD plots along corresponding channels of the linear silicon probe. Passive volume conduction strongly influences the signal recorded at adjacent electrodes in the field potential plots. Some examples are indicated by the red/blue boxes. 1: signal artifacts in CA1; 2: signal artifact at the hippocampal fissure; 3: volume conduction from MPP responses that occlude LPP responses; 4: volume conduction from the granule cell layer response occluding hilar responses. b) Representative sink and source signals and their locations from the CSD plots.

**Figure 3S5.**
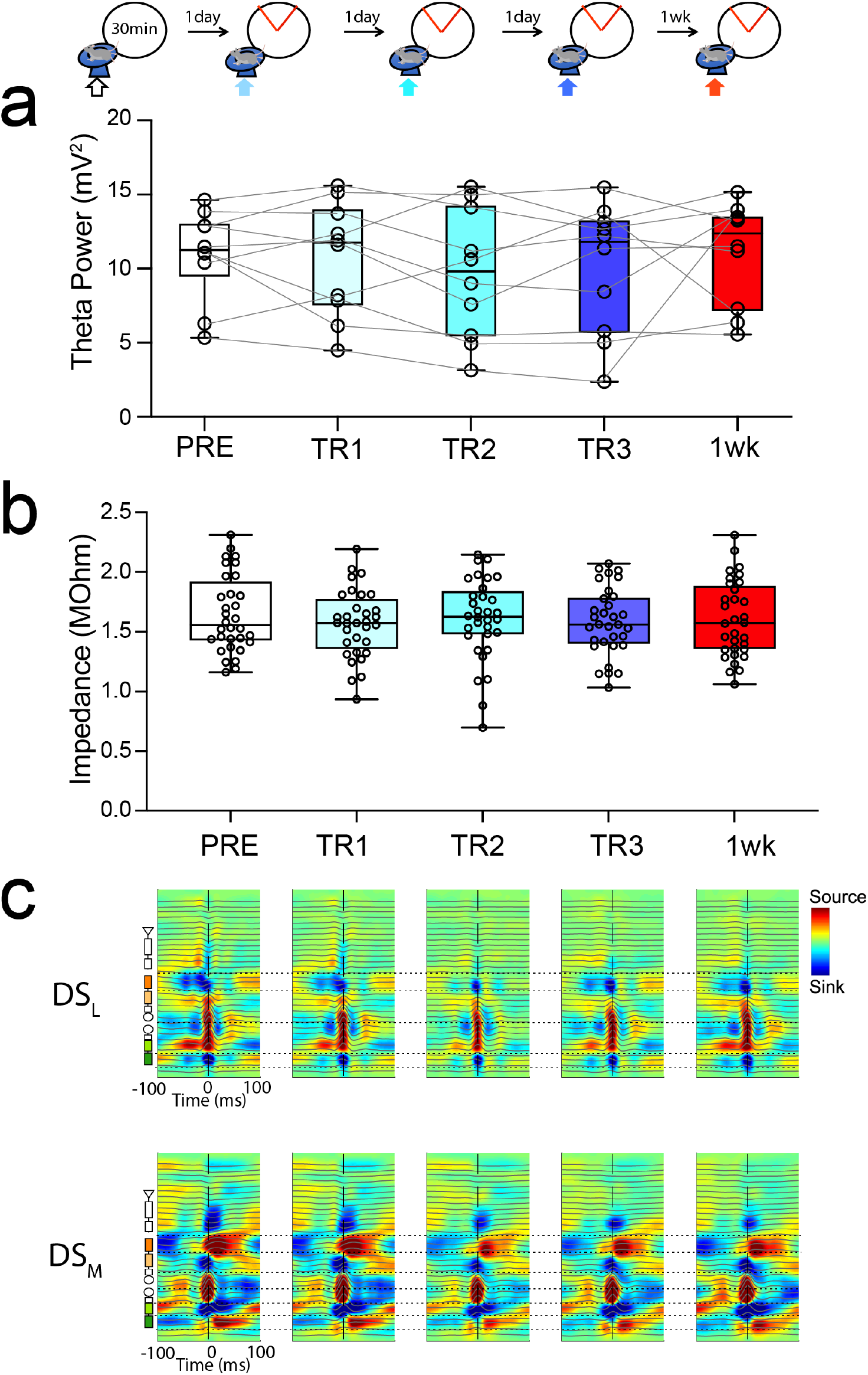
(related to Figure 3). Stability of stimulation and recording electrode properties across the duration of the experiment. a) Average power of spontaneous theta oscillations measured at the molecular-hilar region of supraDG and b) stimulation electrode impedance, both measured during wheel running before active place avoidance training (see schematic at top). Neither theta power (F_2.4, 21.6_ = 0.65, p = 0.55) nor electrode impedance (F_4,128_= 0.294, p = 0.9) changed across the experiment, making it unlikely that changed electrode properties account for the CCT-induced differences in evoked responses. c) Session-averaged current source density (CSD) of LPP-triggered DS_L_ (type 1) and MPP-triggered DS_M_ (type2) spontaneous events also illustrate recording mechanical stability across the experiment.

**Figure 3S6.**
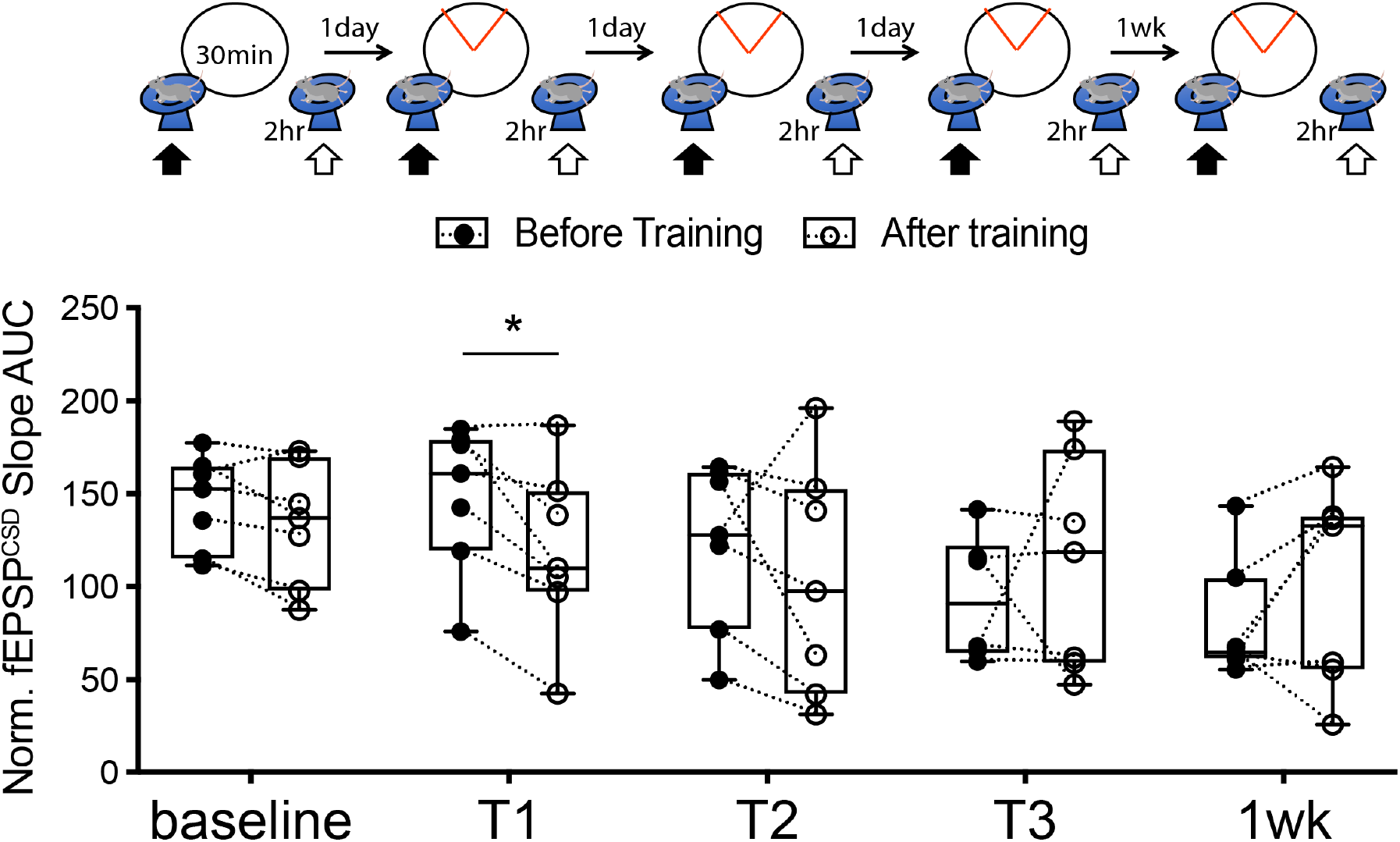
(related to Figure 3). Time course of CCT-induced changes in MPP-DG synaptic responses. fEPSP slope was normalized to the baseline slope prior to exposure to the behavioral arena. The decrease of the fEPSP response is observed within 2 hours of the first CCT trial. * indicates a significant difference from before to after a CCT trial (n = 7, Paired t-test: t_6_ = 3.40, p = 0.01, d = 1.28)

**Fig. 3S7.**
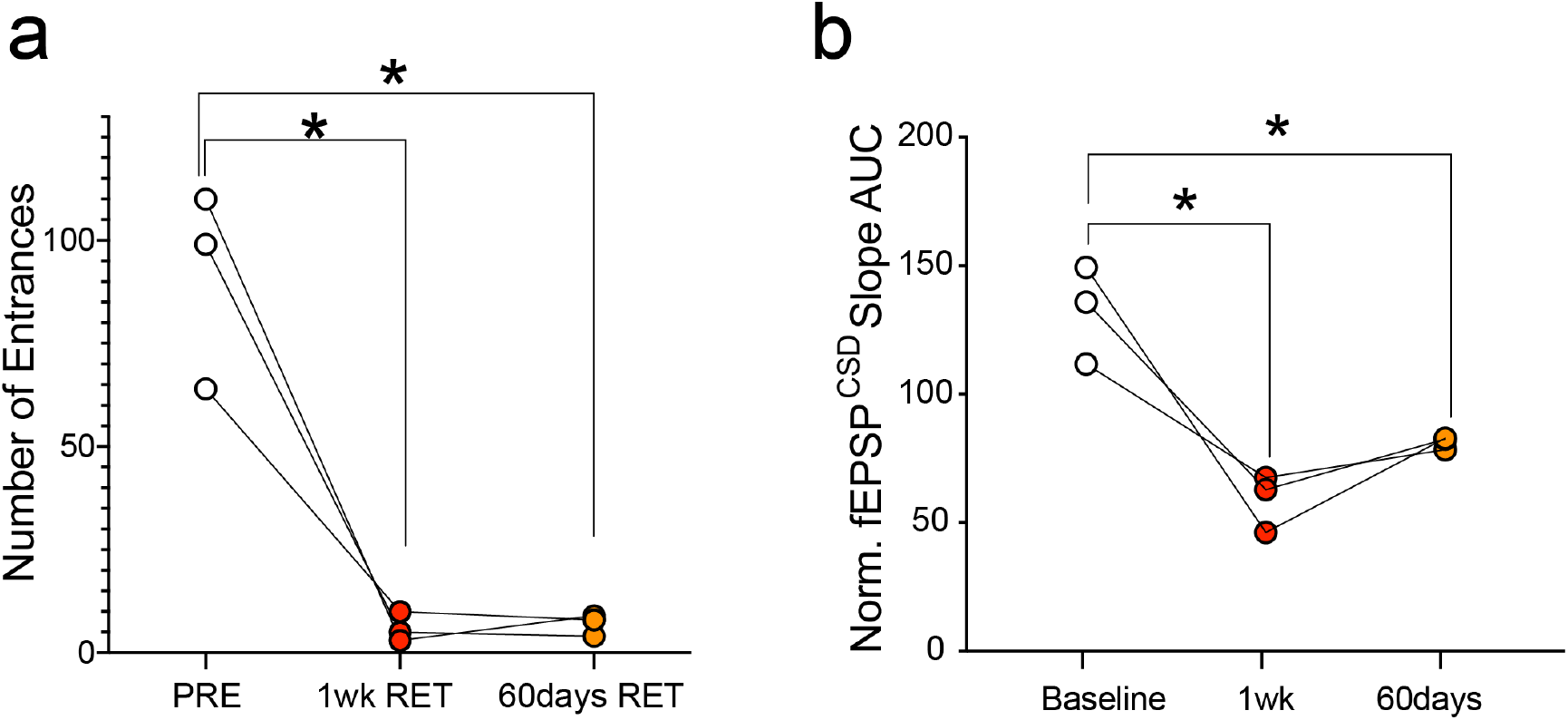
CCT causes place avoidance and reduced population synaptic responses to MPP stimulation that persist at least 60 days. a) Conditioned place avoidance (n = 3, F_1.109,2.038_ = 31.07, p = 0.03, η^2^ = 0.94, pretraining > 1-wk retention = 60-d retention and b) population synaptic responses (F_1.015,2.030_ = 19.30, p = 0.05, η^2^ = 0.90, baseline > 1-wk = 60-d).

**Figure 3S8.**
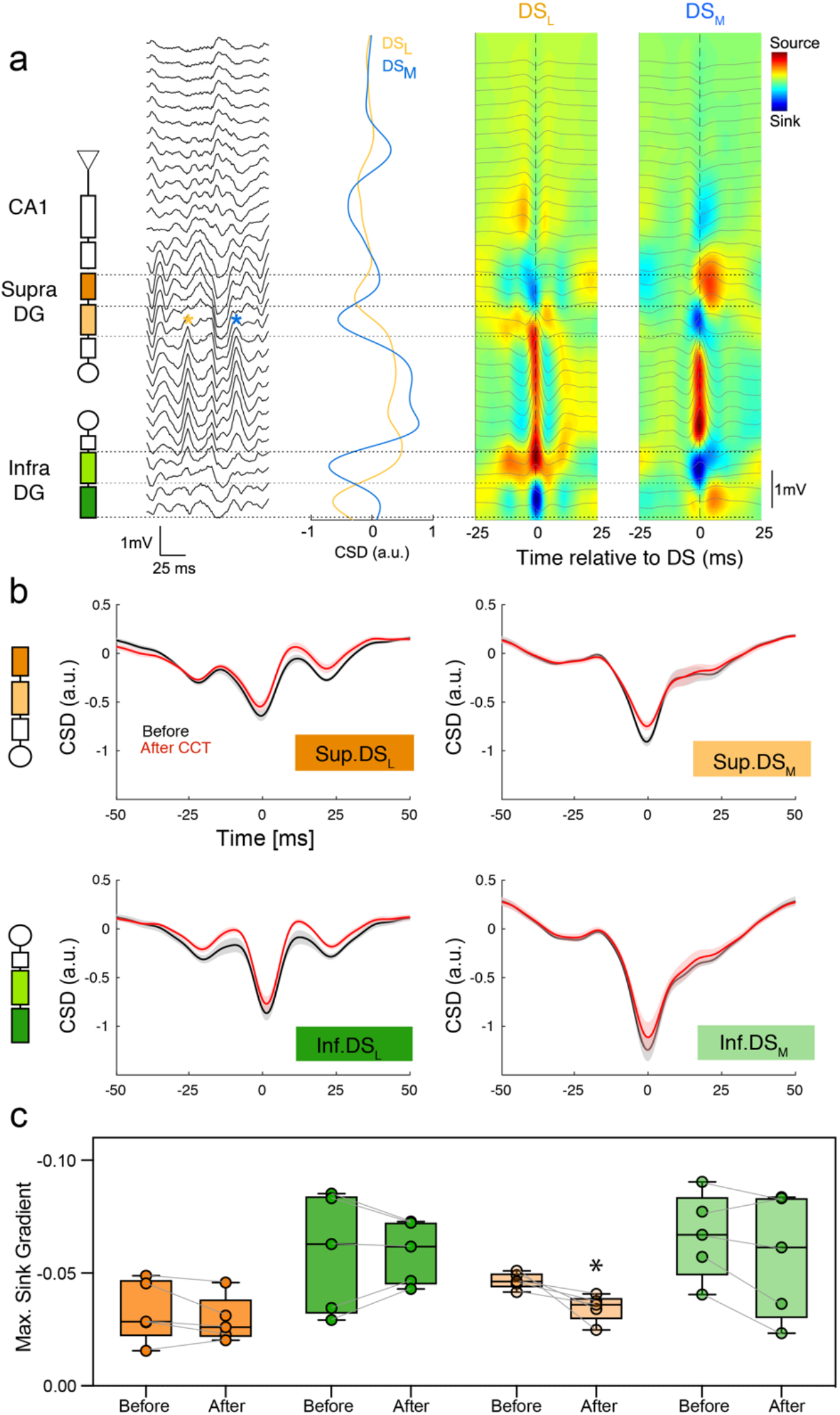
(related to Figure 3). Naturally occurring, spontaneous dentate spikes originating from MPP activation confirm the findings using evoked responses to MPP stimulation. a) A dentate spike (DS, stars) is the DG response to synchronous input from the medial and lateral perforant path, making it a physiological estimate of the MPP-DG connection that is independent of artificial stimulation. DS were identified in the DG hilus as sharp, positive waves of the local field potential with prominence (distance between peak and closest preceding or following trough) greater than 2 S.D. of all detected positive peaks, as well as width between 2.5 and 12.5 ms measured at 50% of the peak’s prominence. b) Two types of dentate spikes – DS_L_ (type 1) and DS_M_ (type 2) were identified using a CSD fingerprinting method, where all DS that exhibited a symmetric pair of current sinks in the distal molecular layers of DG were identified as DS_L_ and DS that exhibited a symmetric pair of current sinks in the medial/proximal molecular layers of DG were identified as DS_M_. The average CSD profiles of DS_L_ (left) and DS_M_ (right) computed from all classified DS events with clearly distinct pairs of current sinks in the distal and medial/proximal molecular layers of DG respectively. c) Average CSD profiles of DS-associated current sinks in molecular layers of DG in supraDG (top row) and infraDG (bottom row) triggered by DS_L_ (left column) and DS_M_ (right column). Black and red colors represent before and after CCT training, respectively. d) Summary comparisons of maximum gradient of the DS sink (i.e. negative slope) measured before and after CCT. CCT only changed DS_M_ (type 2) at the supraDG site but not at the infraDG site, confirming findings assessed by stimulus evoked responses (Paired t-test: t_4_ = 3.04, *p = 0.04, d = 1.36).

**Figure 3S9.**
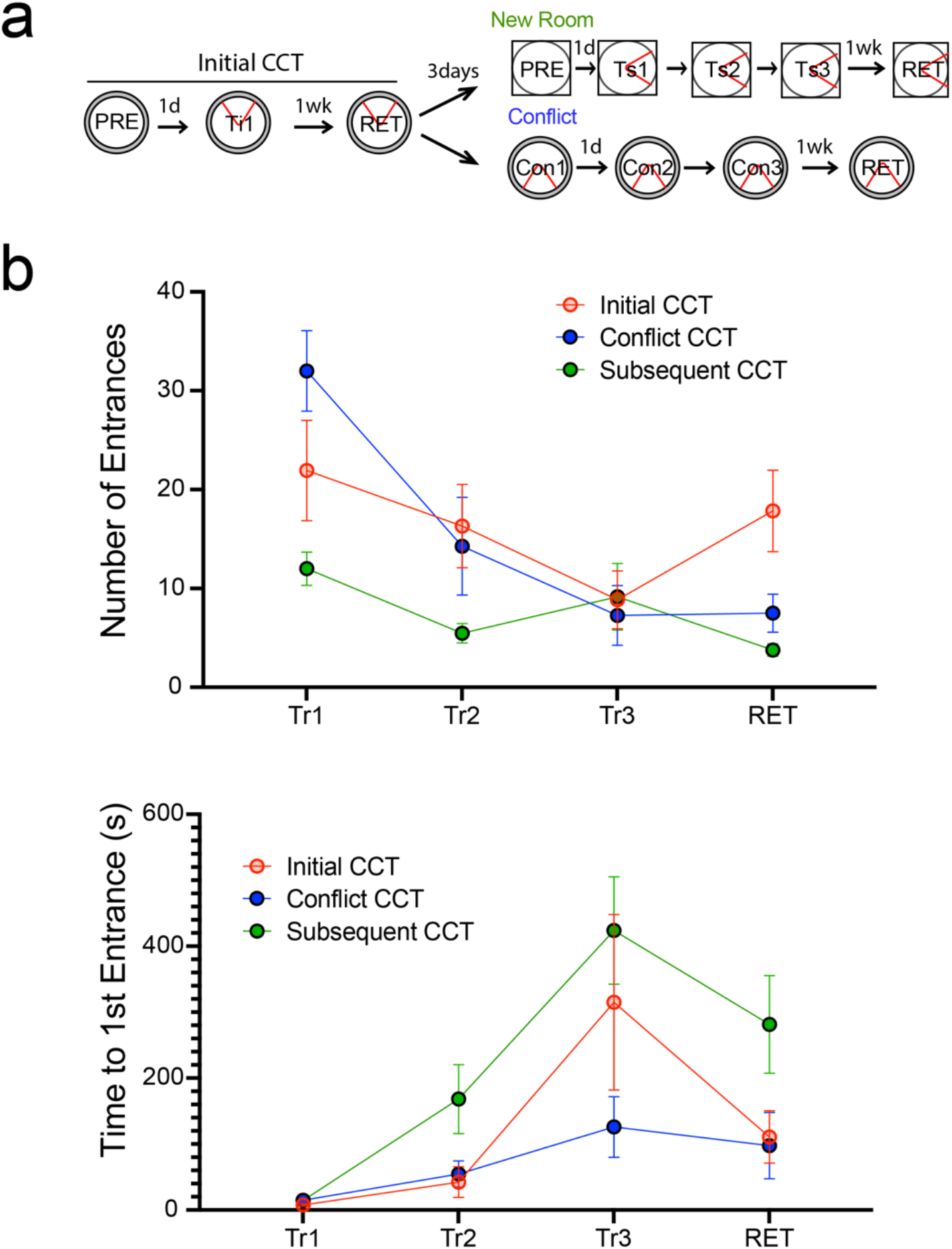
(related to Figure 3). a) Experimental design, b) Learning curves comparing initial CCT training, and subsequent CCT training to either a conflicting location of shock (Conflict CCT), or to a novel location of shock in a novel room (subsequent CCT). (Number of Entrances: Group X Trial RM ANOVA Group: F_2,34_ = 2.95; p = 0.07; Trial: F_3,99_ = 22.56, p = 10^−5^, η^2^ = 0.41; Interaction; F_6,99_ = 5.98, p = 10^−5^, η^2^ = 0.27, Time to 1^st^ Entrance: Group X Trial RM ANOVA Group: : F_2,34_ = 4.42, p = 0.02, η^2^ = 0.21 Trial: F_3,99_ = 13.57; p = 10^−5^, η^2^ = 0.29; Interaction; F_6,99_ = 1.581, p = 0.16).

### Supplemental Results – Persistent Experience-Dependent Changes in Inhibition

**Figure 4S1.**
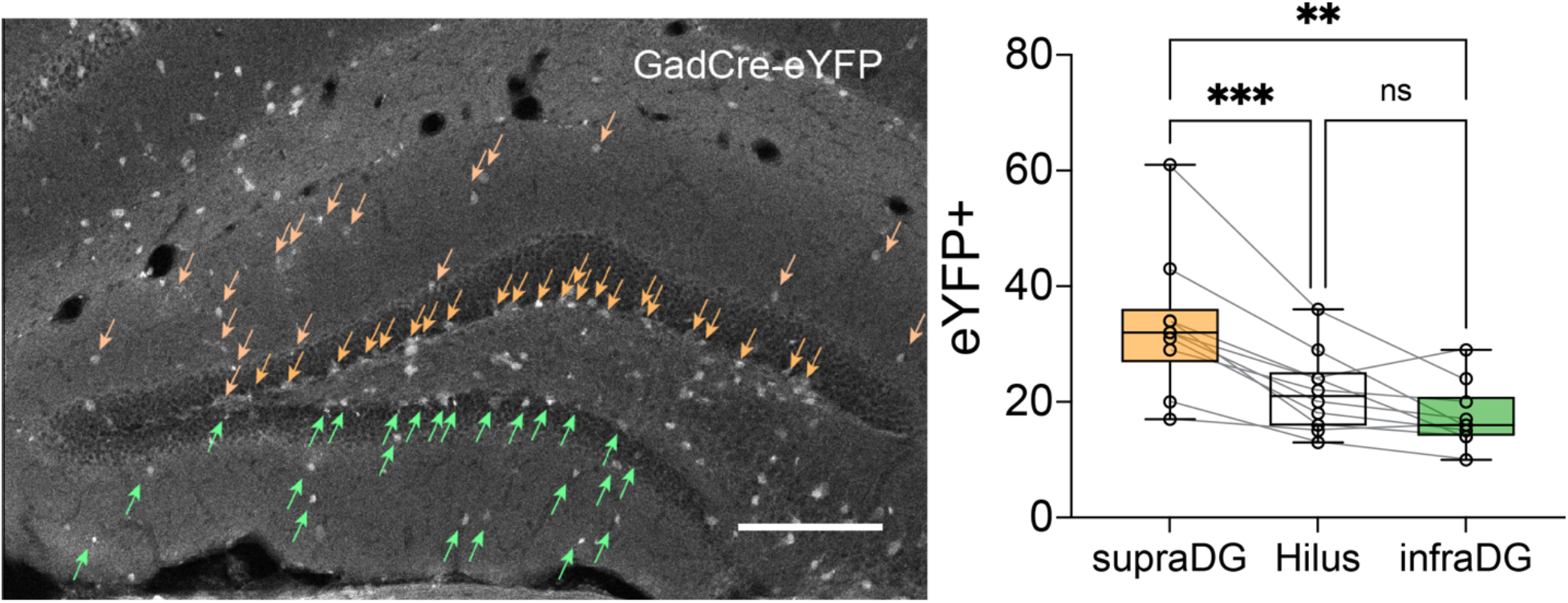
(related to Figure 4). Gad2-expressing interneurons are more abundant in supraDG than infraDG. Left) eYFP expressing interneurons in Gad2Cre-eYFP mice (scale bar 0.2 mm); Right) cell counts per section in mice after CCT (n=6) or SE (n=4) experience document enrichment of inhibitory interneurons in supraDG relative to infraDG (F_1.215,10.93_ = 21.40, p < 0.001, η^2^ = 0.69, SupraDG > InfraDG = Hilus). Orange and green arrows indicate eYFP+ cells in the supraDG and infraDG, respectively.

**Figure 4S2.**
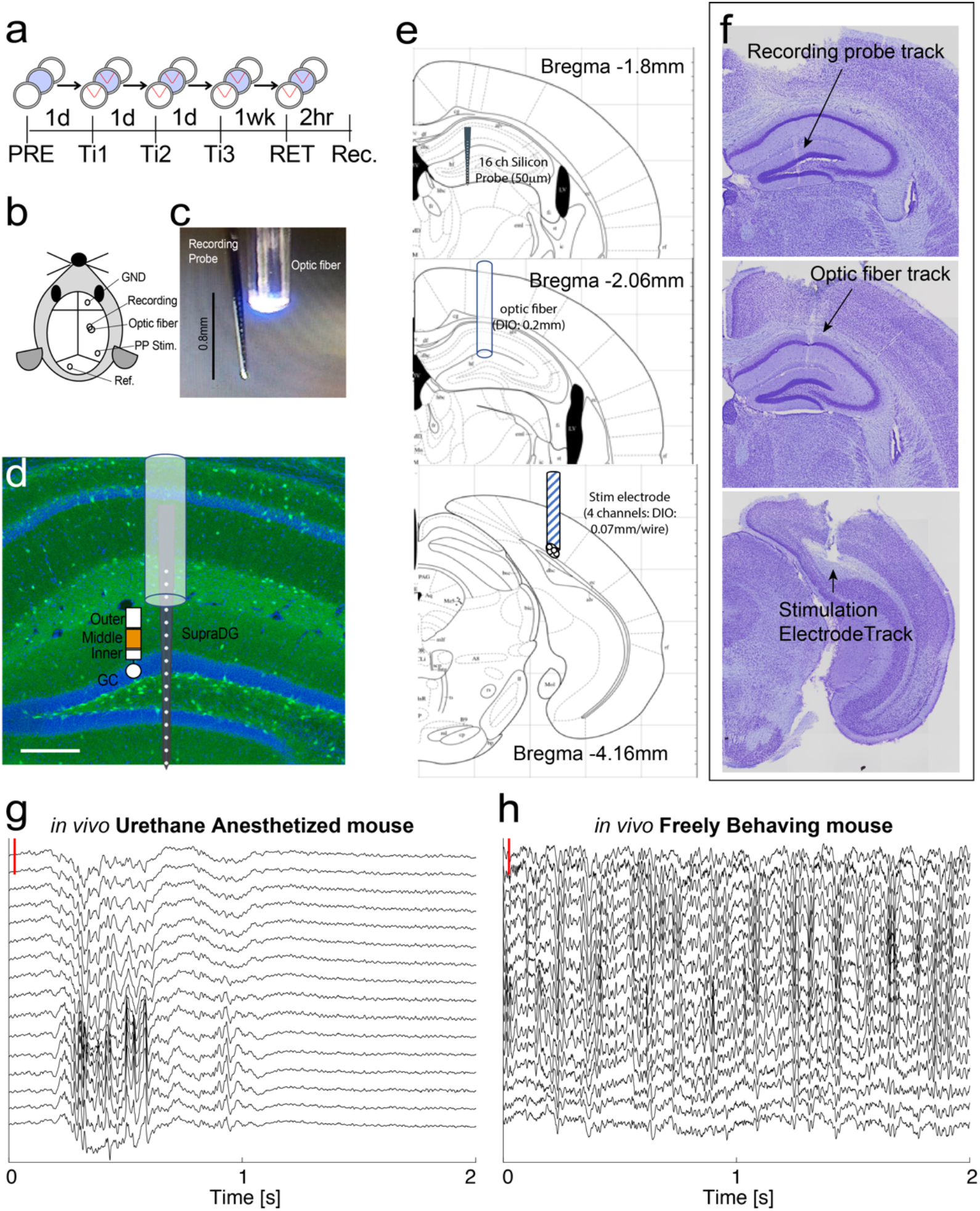
(related to Figure 4). The *in vivo* experiment under urethane anesthesia to investigate the role of inhibition in the CCT-induced effects. a) Experimental design, b) schematic of the preparation, c) photo illustrating the recording electrodes and optical stimulation, and d) The schematic is superimposed on a histological section of a Gad2-Cre-eYFP mouse’s dorsal hippocampus with eYFP (green) and immunofluorescence for DAPI (blue), to indicate the circuit that was targeted (scale bar 0.02mm). e) Atlas from Franklin and Paxinos^47^ and f) corresponding histological sections illustrating electrode and optical fiber placements. Sixteen-channel linear electrode array recordings of LFPs across the somatodendritic axis of dorsal hippocampus g) under urethane anesthesia and h) in the freely-behaving mouse. Under urethane, LFPs reflecting rhythmic ongoing synaptic activity is much attenuated compared to in the behaving mouse. Voltage scale (red bar) = 1 mV.

**Figure 4S3.**
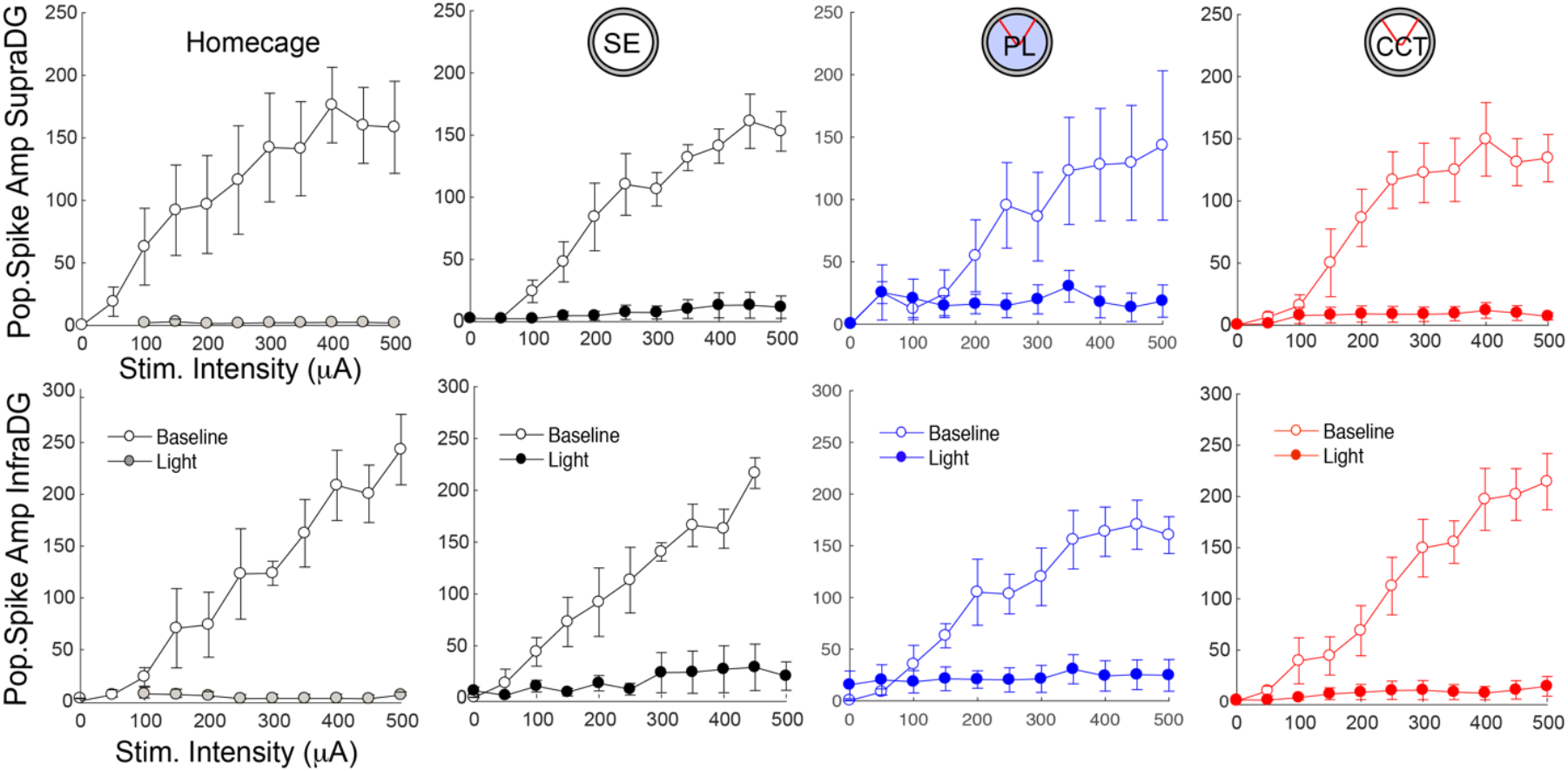
(related to Fig 4). Population spike responses are blocked by activating ChR2 in Gad2-expressing cells 5 ms before MPP stimulation *in vivo* under urethane anesthesia, regardless of prior training. Input-output relationships of the response to MPP stimulation with and without prior light activation of inhibitory-neuron ChR2 in the (upper) supraDG and (lower) infraDG. This demonstrates light-stimulated inhibition is effective in all groups.

**Figure 4S4.**
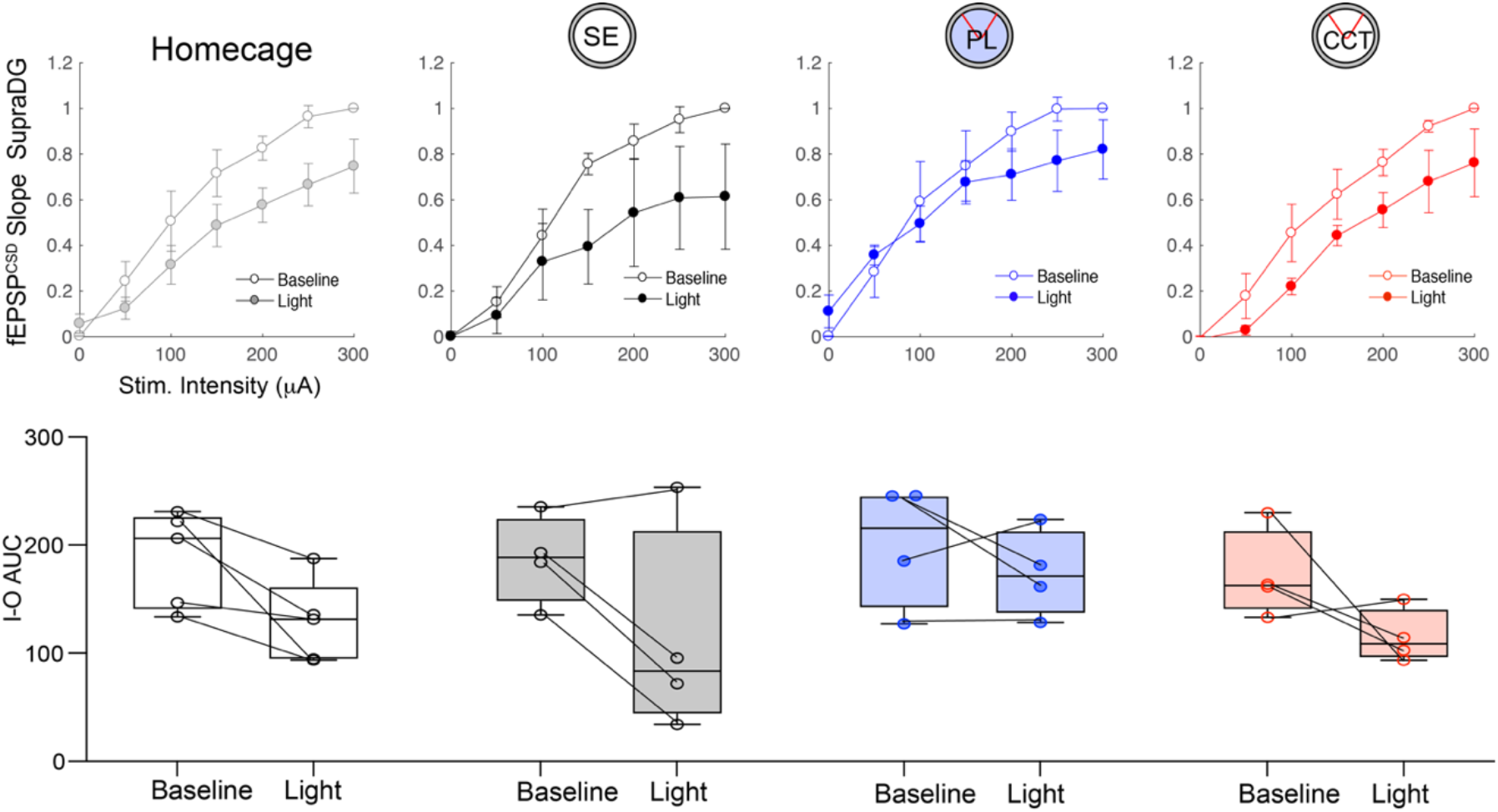
(related to Fig 4). fEPSP evoked responses at supraDG are reduced by activating Gad2-expressing cells at the same time as MPP stimulation *in vivo*, regardless of prior training. Upper) Input-output relationships of the response to MPP stimulation with and without concurrent light activation of inhibitory-neuron ChR2. Lower) The corresponding summaries represented as the area-under-the-curve (AUC) for statistical analysis of the group and light stimulation effects. Group X Stimulation ANOVA, Group: F_3,13_ = 0.80, p = 0.51; Stimulation: F_1,13_ = 16.05, p = 0.0015, η^2^ = 0.55; interaction: F_2,10_ = 0.4921, p = 0.70).

**Figure 4S5.**
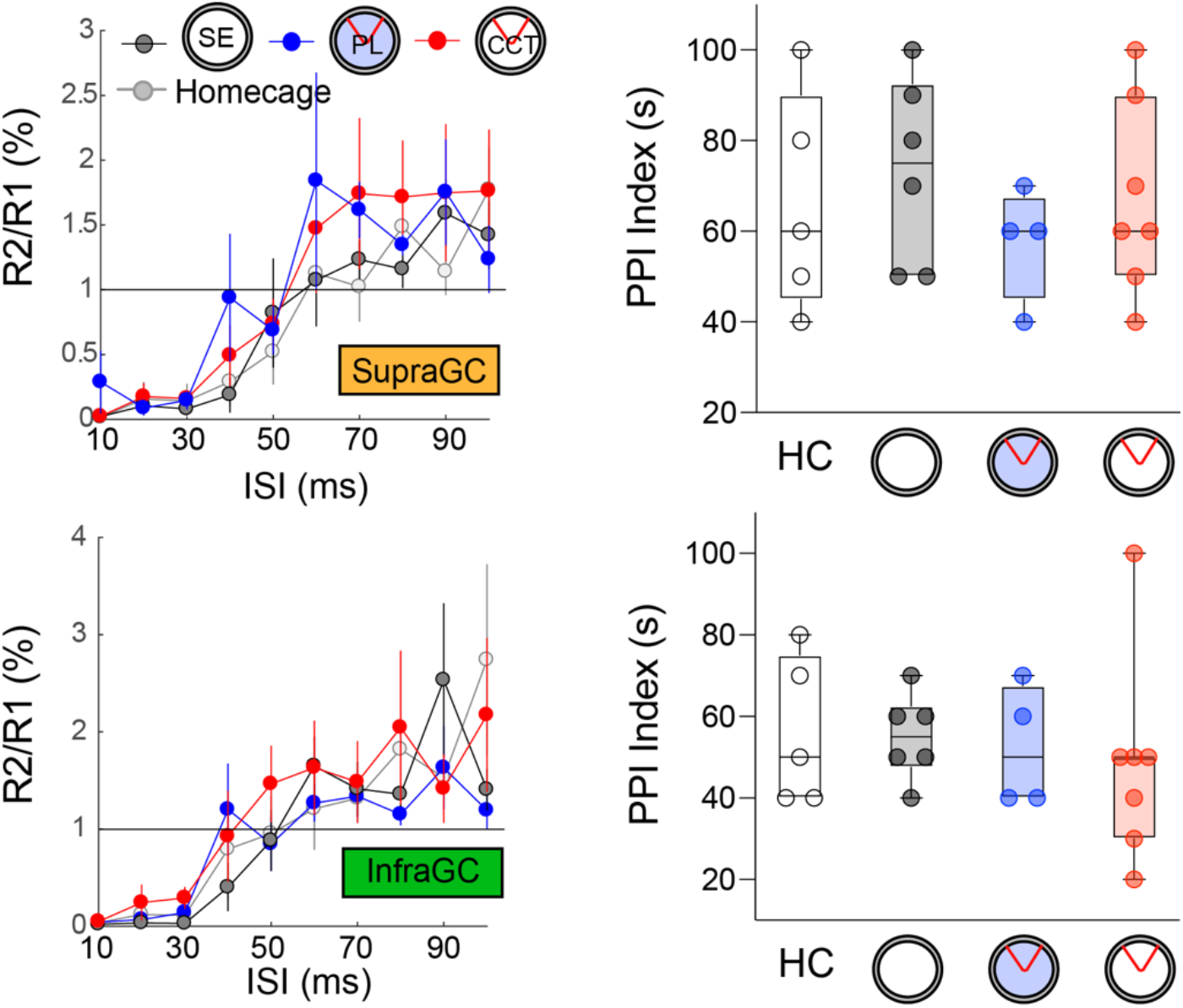
(related to Figure 4). Paired-pulse inhibition and facilitation measured at the granule cell layer of DG are not changed in CCT mice. Responses to MPP stimulation were recorded in urethane-anesthetized mice at the supra and infra granule cell layers of the DG. Significant changes were not detected although there is a hint of enhanced facilitation in the CCT group. (supraDG granule cell Group: F_3,18_ = 0.47, p = 0.70; InfraDG granule cell group: F_3,18_ = 0.19, p = 0.90).

**Figure 4S6.**
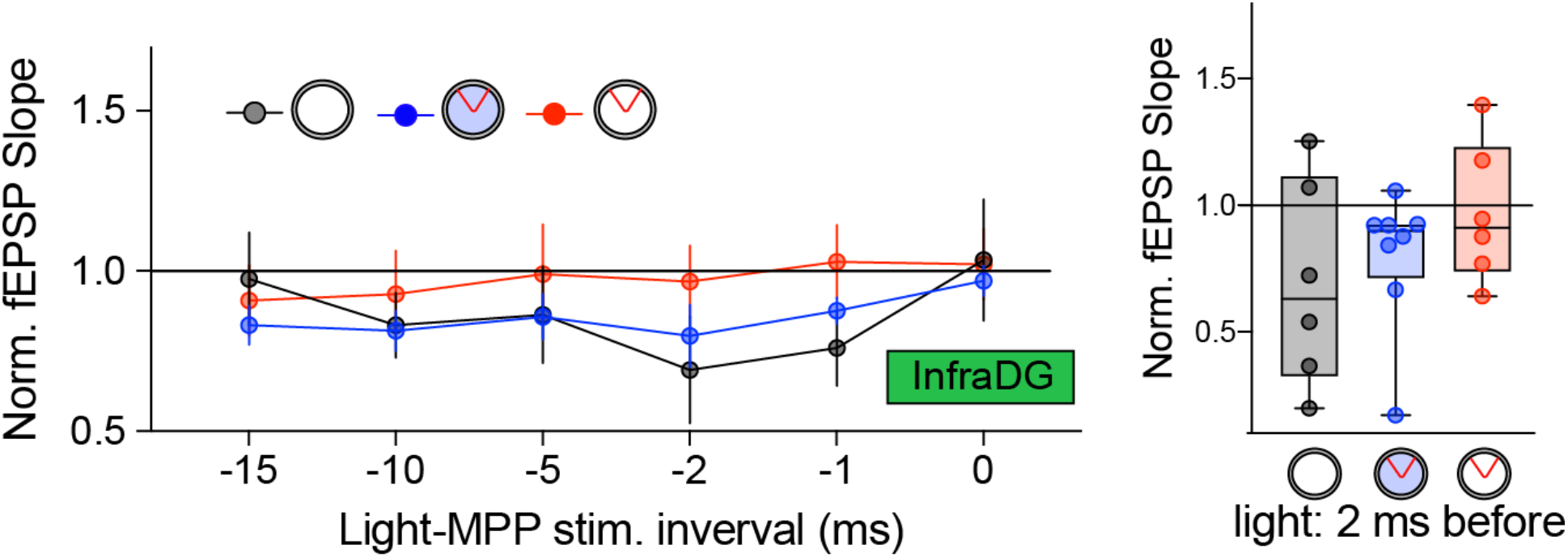
(related to Fig 4). Response recorded in infraDG to MPP stimulation with ChR2 activation of inhibitory neurons at different time offsets. left: infraDG (2-way RM ANOVA Group: F_2,17_ = 0.50, p = 0.62, Light-Stim ISI F_2.06,34.95_ = 2.82, p = 0.73; interaction: F_10,85_ = 1.15, p = 0.34) right: Light 2ms before (F_2,17_ = 1.14, p = 0.34).

**Figure 4S7.**
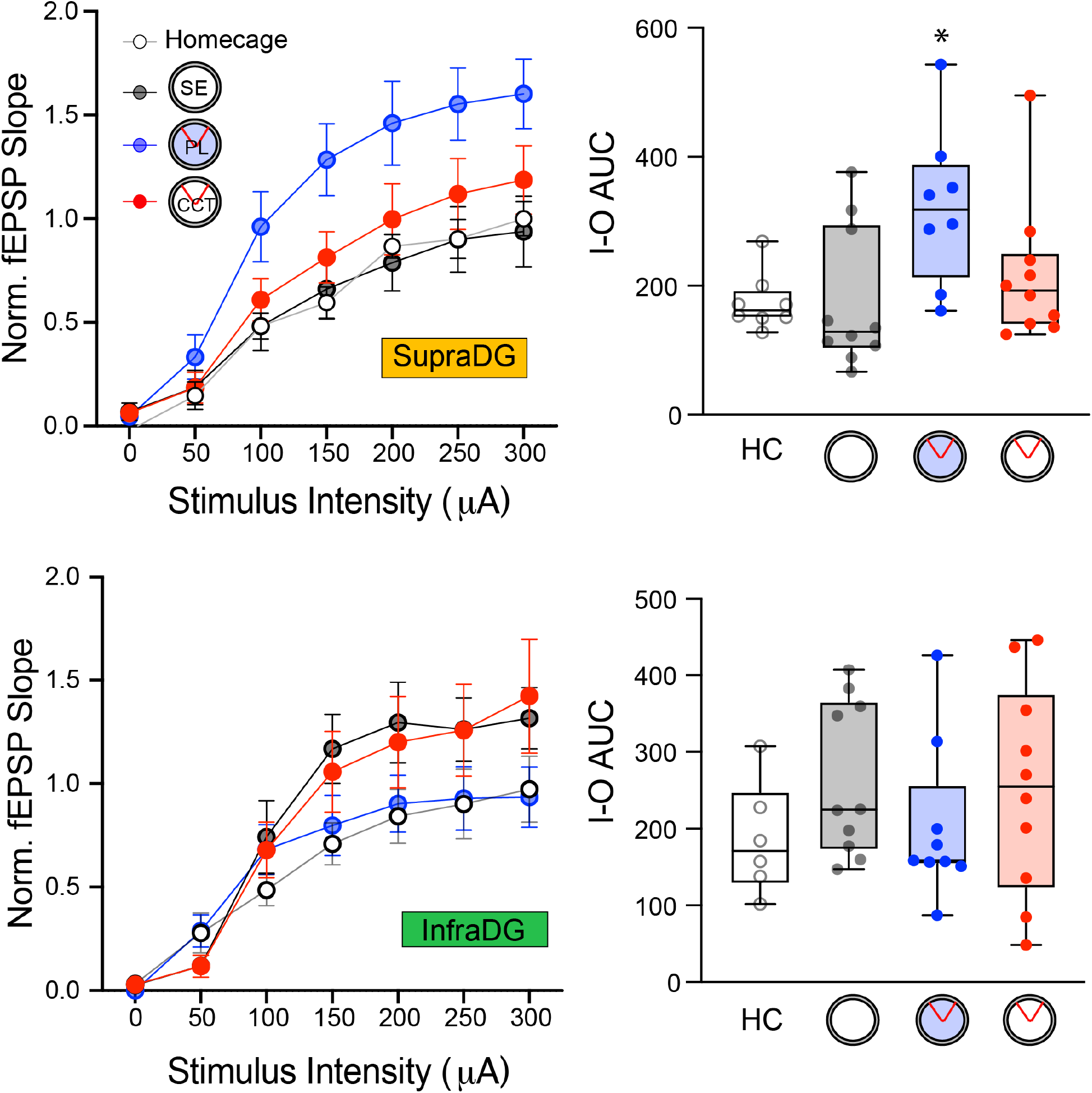
(related to Fig 4). Input-output relationship of the response to MPP stimulation in *ex vivo* hippocampal slices. I-O curves (left) and their summary as the area-under-the-curve (AUC, right) upon which statistical group comparisons were made. (Upper) SupraDG: IO-AUC F_3,32_ = 3.80, p = 0.02, η^2^ = 0.26, Sidak’s multiple comparisons: HC vs PL p = 0.021. (Lower) InfraDG: IO-AUC F_3,31_ = 0.93, p = 0.44

## SUPPLEMENTARY INFORMATION

### Ethics Approval

All work with mice was approved by the New York University Animal Welfare Committee (UAWC) Protocol ID: 15-1453.

### Active Place Avoidance Training

The mice (C57BL/6J; n = 42) were placed inside the rotating arena on which they are confined by a 40-cm diameter, 50-cm high transparent cylindrical PETG plastic wall. Some of these animals had been implanted so that the response to medial perforant path (MPP) stimulation could be assessed across their training. The behavior of 39 animals was analyzed, and three were not because the electrophysiological signal quality was not sufficiently maintained across the training protocol.

The electrifiable floor was either made of parallel stainless steel rods or copper mesh so that a 500-ms, 60-Hz 0.2-0.3 mA constant current foot shock could be delivered to the mouse. On the copper mesh floor, the shock was delivered across the high impedance of the mouse’s paws on the mesh and via a counter-balanced cable, a low-impedance subcutaneous shock electrode at the neck. The shock was scrambled across the parallel rod floor by switching across 5 poles to ensure the mouse could not avoid shock by assuming a particular posture, and that shock could not be shorted by feces. Windows-based software (Tracker 2.36, Bio-Signal Group Corp., Acton, MA) determined the mouse’s location in every video frame from an overhead video camera. On the mesh floor, the mouse’s position was determined by tracking the location of an infrared LED on the clip that connected the cable to the subcutaneous shock electrode. On the parallel rod floor, the software tracked the centroid of the mouse’s body in the video image. The software-defined a 60° Room-frame shock zone that was at a fixed location in the room. During training sessions, the computer turned on shock whenever the mouse entered and remained in the shock zone for 0.5 s. The shock was repeated every 1.5 s until the mouse left the shock zone. The software automatically controlled rotation of the arena at 1 rpm as well as delivery of shock. End-point measures were computed using TrackAnalysis 2.5 Bio-Signal Group Corp., Acton, MA).

### Experimental design: Active place avoidance

The behavioral protocol was optimized to evaluate acquisition of learning to learn as well as for collecting sufficient electrophysiological data before, during, and after the cognitive control training. There were two training phases, the “Initial Training” and “Subsequent Training” phases. Each daily session consisted of a 30-min acclimatization to the experimental room, a 30-min session on the behavioral arena, and 2 hours post-training in the room. The first day, the subject was transferred to the experimental room to acclimate for 30 min. Then there was a pretraining session on the rotating arena with the lights on and no shock. The animals were free to explore the experimental apparatus without any reinforcement. After pretraining, the mouse was returned to its home cage and stayed in the experimental room for 2 hours. The mouse was then transferred to the vivarium. Twenty-four hours later, the mouse was returned to the experimental room for the 30-min acclimatization, followed by a training session on the rotating arena. The only difference from pretraining is that on training trials the mouse received a foot shock whenever it entered the shock zone. The same training sessions were repeated on the next two days, for a total of 3 training sessions. After a week in the vivarium, the mice were returned to the experimental room to test for retention of the conditioned avoidance. The experimental conditions were identical to the training sessions.

Three days after retention of the conditioned avoidance was evaluated, the mice began the Subsequent Training phase. A different experimental room was used; it was located on the opposite side of the lab and had substantially different spatial cues, a different type of arena flooring (mesh or rods), and a different ambient odor (neutral or vinegar). The behavioral protocol during Subsequent Training was identical to the Initial Training phase: one day of pretraining without shock, training for three days followed by a 1-week retention test.

### Experimental groups

Three groups of mice were used to investigate whether Cognitive Control Training (CCT) specifically causes learning to learn. In addition to the CCT group, there were also Place Learning (PL) and Spatial Exploration (SE) control groups. The CCT group received R+A-training during the Initial Training phase. The R+A-notation indicates avoidance of a room place was conditioned and arena places were not conditioned. The PL group received R+ training during Initial Training on the rotating arena filled with ~1 cm deep 1% NaCl solution, which removed Arena-frame odor cues but allowed the mice to receive foot shock. The R+ notation indicates that avoidance of a room place was conditioned and that arena locations were not salient. The SE group never experienced shock during the Initial Training phase but other than shock, they experienced the identical physical environment as mice in the CCT group. Mice in all three groups had R+A-CCT for the Subsequent training phase.

### T-maze

A T-maze was used to evaluate whether the learning to learn that is caused by CCT generalizes to different tasks that require cognitive control. The T-maze had transparent walls made of 0.8 cm thick acrylic. The width, the length and the height of the maze walls were all 40 cm. The floor was made of parallel rods that could be electrified, scrambled across 5 poles. At the beginning of the task, the subject was placed in the start arm and allowed to explore the maze without shock for 10 minutes before being returned to their home cages. Fifteen minutes afterwards, T-maze training began. Each of the three sessions had 10 trials. The mouse’s location was tracked during maze behavior (Tracker 2.36, Bio-Signal Group Corp., Acton, MA). For each trial, the mouse was placed in the start arm. After 5 seconds 500 ms, 60 Hz, 0.3 mA foot shock was automatically turned on by the tracking software if the mouse was detected in the shock zone, defined as the start arm and the punished arms. Shock was automatically repeated every 2.5 s until the mouse escaped to the safe arm. After it remained in the safe arm for 5 seconds, the experimenter removed the mouse. The inter-trial interval was 10 seconds. After 10 trials, the mice were returned to the home cage for a 10-minute rest. In the second trial, the safe and punished arms were switched and mice had again 10 trials to escape shock. After another 10-min rest in the home cage, the mouse had a third session of 10 trials with the safe and punished arms switched again. Successful avoidance was scored when the mouse avoided shock by escaping to the safe arm within 5 s. We scored the T-maze behavior by counting the number of trials to reach a success criterion. The criterion was set to be the second trial when the mouse escaped to the safe arm and remained there to successfully avoid the shock. After meeting this criterion, mice tended not to make any further errors.

### Cannulation surgery and intracranial muscimol infusion

Subjects were 10 adult C57BL/6J male mice weighing approximately 30 g. Animals were housed individually in plastic cages with free access to water and mouse chow. The colony room was maintained under a 12:12 h light/dark cycle (lights on at 7:00 a.m.) and all behavioral tests were conducted during the light portion of this cycle. All procedures were approved by the Animal Care and Use Committee at New York University. Subjects were adapted to handling for three to four consecutive days before surgery. Immediately before surgery, each mouse was anesthetized with an intraperitoneal (IP) injection of sodium pentobarbital (50 mg/kg). Animals were then prepared with bilateral stainless steel 31-gauge cannulae aimed at the dorsal hippocampus using stereotaxic coordinates (1.94 mm posterior, ±1 mm lateral, 1 mm ventral relative to Bregma ^1^). Cannulae were secured to the skull with stainless steel screws, Vetbond, and dental cement. Following surgery, subjects were treated with Ketoprofen (5 mg/kg) for 3 days to minimize discomfort during the recovery period. Stainless steel mock injection needles remained in the cannulae when mice were not being injected to prevent occlusion. Each mouse was given a recovery period of at least 5 days before behavioral testing.

After the recovery period, the mice went through an active place avoidance training protocol with 30-min training sessions and 1-day inter-training intervals. A week after the last training session, there was a 5-minute retention session with shock off. A day after the retention test, the mouse received a 30-minute retraining session in case there was extinction learning during the retention period. Five days after retraining, either muscimol or vehicle (PBS, pH 7.4) was infused into the hippocampus. Thirty minutes after the infusion, mice were transported to the behavioral testing room and given a 5 min retention test with shock off. After the retention test, mice were returned to their home cages for 24 hours after which they received a second retention test. All subjects were habituated to the injection procedure for the three days preceding infusion. Each mouse was wrapped in an absorbent pad, and gently restrained by hand while the infusion pump was activated. Drugs were prepared fresh on the day of infusion. Muscimol was dissolved in sterile saline (pH ~7.0, Teknova, Hollister, CA) to create a final concentration of 10 mM. All subjects received an infusion volume of 0.5μl (injection speed: 0.1μl/min) aimed at the dentate gyrus of the dorsal hippocampus. The injection cannulae, which were cut to extend 0.5–0.7 mm beyond the guide cannulae, remained in place for an additional 5 minutes after the infusion. The subjects were returned to their home cages immediately after the infusions.

### Electrode implant surgery and electrode location confirmation

Subjects were 20 adult C57BL/6J male mice weighing approximately 30 g. Mice were anesthetized with 1.75% isoflurane and placed in a Kopf stereotaxic frame to implant three bone screws, a recording electrode and a stimulation electrode. A 16- or 32-site linear silicon probe arrays with 30-μm diameter recording sites and 50-μm inter-site spacing (Neuronexus, Ann Arbor, Ml; p/n: A1×16-3mm-50-703 or A1×32-6mm-50-703) was placed in the dorsal hippocampus (2 mm posterior, 1.0 mm lateral; −1.8 ~ 2.3 mm ventral from Bregma) to span all layers of DG. A four-wire stimulating electrode bundle was made by twisting together four 75-μm diameter nichrome wires (California Fine Wire, Grover Beach, CA). The bundle was cut at an angle so as to span 0.5 mm. During surgery, the stimulating bundle was placed in the ipsilateral perforant path (0.5 mm anterior, ±4.1 mm lateral, 1.0-1.6 mm ventral from lambda).

The stimulation electrodes were each attached to a pin in a Mill-Max connector. The two-wire combination that evoked the largest response at the recording electrode was selected for the stimulation experiments. During surgery, the location of both electrodes were monitored and verified by layer-specific spontaneous activity as well as layer-specific evoked potential waveforms. Typically, the largest theta amplitude was observed in the DG. Dentate spikes were used as an indicator of the DG location. The tip of the recording electrode was placed at the ventral limit of the DG. After confirming the placement of the recording electrode, the stimulation electrode was lowered down while 200-300 μA constant current test pulses were delivered at 15 s intervals. Evoked potentials have different characteristic waveforms at different stimulation locations, which served as a further guide and verification of electrode placement. Evoked response waveforms were carefully checked with different combinations of stimulation electrode channels. After confirming that the recording electrode array extended through CA1 and DG and that the stimulation electrode was in the medial perforant path, the wires and connectors were fixed in place with dental cement. The Omnetics connector of the recording electrode and the Mill-Max connector of the stimulating bundle were anchored to the skull along with bone screws using one of two dental cements (Grip Cement, Dentsply, Milford DE and TEETs Denture Material, Cooralite Dental Mfg, Diamond Springs, CA).

A constant current stimulus isolation unit (WPI, Sarasota, FL; model: A365RC) was used to deliver individual unipolar 100 μs stimulus pulses across the electrode pair. A customized amplifier and digitization board with 32 unipolar inputs ^2^ (RHD2132, Intan Technologies, Los Angeles, CA) was connected directly to the recording electrode assembly for signal amplification and digitization. A lightweight cable (Intan Technologies, Los Angeles, CA) was counter-balanced, and used to power the amplifier board and an infrared LED that was used to track the mouse’s position. This cable also transmitted digital data to the computer using a custom recording system that was connected to the USB port of a PC. The cable was connected through a lightweight swivel to enable free movement of the animal. Signals from each electrode were low-pass filtered (500 Hz) and digitized at 2 kHz for local field potentials (LFPs) and low-pass filtered (4000 Hz) and digitized at 8.12 kHz for evoked responses. The LFP data were first processed by a custom artifact rejection algorithm, which marked continuous segments of artifact-free signals. Such segments that were 4 s or longer were used for further analysis. Signal artifacts were defined as signals crossing a predefined threshold selected by visual inspection and used for analysis of the entire dataset. Each artifact segment was extended by 0.25 s on both sides and all artifact-free segments smaller than a 1-s window were also discarded. Each channel in the dataset was processed independently and the algorithm performance was verified by visual inspection. Three animals were excluded because the electrophysiological signal quality was not sufficiently maintained across the training protocol.

### Histology

Upon conclusion of the experiments, all mice were transcardially perfused with 1X phosphate buffered saline (PBS) followed by 4% PFA. The brains were extracted and stored in formalin overnight. The brains were stored in 30% sucrose in 1X PBS until they were cut on a cryostat (40 μm) and thaw mounted onto gelatin-coated slides. The sections were dried overnight at room temperature and then Nissl stained. The slides were scanned at 10X with an Olympus VS120 microscope and the images were subsequently examined for electrode tracks to verify the stimulation and recording locations.

### Behavior Training and Evoked potential recording

After the electrode implant surgeries, animal subjects had 7-10 days to recover. The animals were handled to get comfortable with the experimenters, cable connections, and transportation to and from the testing rooms for 3-5 days. During this acclimatization period, the mice received one 10-min electrode screening session to examine the recording quality. If there were significant reductions of theta amplitude, electrode location changes greater than 5 channels (250 μm), or weak or no evoked potential responses, the mice were excluded from the experiment (n = 3). On the training day, mice in their home cages were moved to the behavior testing room to habituate to the room for 30 min. During the habituation period, LFPs were recorded and used as a recording for each subsequent recording session. For evoked potential recording sessions, the mouse in its home cage, was confined on a plastic horizontal running wheel by a clear plastic cylindrical wall. The stimulation electrode was connected to deliver test pulses that varied in intensity from 0 to 250 μA. The test pulses were given to measure the evoked potential responses in the dorsal hippocampus. Each pulse intensity was repeated 5 times at 15-s intervals so as to collect the data for constructing an input (I, stimulus intensity) - output (O, evoked response) I-O curve that evaluates the functional MPP-DG connection. The mice were running continuously on the wheel during the I-O data collection and the movement was monitored using video tracking, which ensured that the mice were in a behavioral and electrophysiological steady state characterized by running-associated theta oscillations in the hippocampal LFP.

Behavioral training sessions were 30 min long without shock (Pretraining or SE group) or with 0.2-0.3 mA location-specific foot shock conditioning (training sessions for the CCT and PL groups). Right after the training session, the mice were returned to their home cages in the experimental room and allowed to rest for 2 hours. Afterwards, one more post-training I-O recording session was conducted under identical conditions as the pretraining session. The experimental procedures were identical for all three groups, except for the absence/presence of shock conditioning during training (SE vs. CCT and PL) and the arena floor (shallow 1% NaCI solution on the arena for the PL group, that was 1-2 cm deep sufficient to just cover the floor rods). Importantly, all physiological measurements were conducted during running on the wheel, in identical physical and behavioral conditions for all mice.

### Current Source Density (CSD) analysis to localize fEPSP changes in linear multi-channel array recordings

CSD analysis was performed to attenuate effects of volume conduction on each of the layer-specific responses. The CSD is computed as the second spatial derivative of the voltage along the recording depth:

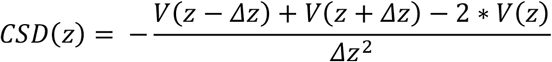

where *CSD*(*z*) is the CSD at depth *z*, *V*(*z*) is the Voltage at depth *z* and Δ*z* = 50 μm is the distance between adjacent recording sites. The CSD is calculated in units of mV/mm^2^ and can be multiplied by a conductivity constant ∫ = 0.3 S/m resulting in units μA/mm^3^. By convention, CSD is computed as the negative of the second spatial derivative with negative CSD indicating an extracellular current sink and positive CSD indicating an extracellular current source. Because it is isotropic ^3^, we assume that the conductivity is homogeneous within the hippocampus as it was shown that variations in the conductivity had little effect on hippocampus CSD estimates ^4^. The CSD is numerically computed using the iCSD toolbox with spline fitting ^5^ where the conductivity is set to be equal among tests.

The CSD was used to locate the extracellular current sinks and sources after perforant path stimulation and in so doing localize the electrode to specific anatomical sites for each recording, independently of other recordings. From perforant path stimulation, the granule cell layer was identified by the largest source in the lower channels that also included a sink, which was the population spike activity. The hilar responses were identified as a decreased source between the suprapyramidal and infrapyramidal granule cell layers. From the granule cell layer, the molecular layer was identified as the largest sink 1-2 channels (50-100 μm) above and below the two granule cell layers. Quantification of evoked potentials from the resulting CSD traces was performed on the physiologically/anatomically localized channels. The molecular layer fEPSP response was taken as the negatively sloped response from the channels at the supra- and infrapyramidal molecular layers. The population spike amplitudes were measured at the supra- and infrapyramidal granule cell layers.

### Evoked potential recording and analysis

Each input pathway’s specific stimulus-response features were measured from input-output (I-O) curves. We analyzed the field excitatory postsynaptic potential (fEPSP) and population spike (PS) responses from the CSD. fEPSP and PS responses were analyzed off-line using custom MATLAB code. The start of fEPSP responses were visually identified as linear decreases in the fEPSP responses 1-2ms after the stimulation was delivered. The maximum decay slope was computed across 0.5 ms windows, and the average of the 5 responses to a specific stimulus intensity was used to characterize the fEPSP slope for the stimulus intensity. The I-O curves were generated from stimulus intensities ranging from 0 to 250 μA in steps of 50 μA. At each stimulus intensity, five responses were recorded with 15 s inter-stimulation intervals.

### In vivo Paired-pulse stimulation

Paired-pulse stimulation to assess paired-pulse inhibition was performed at 70% of the intensity required to elicit the maximal PS response. Stimulus pairs were delivered at varying interstimulus intervals ranging from 10 to 100 ms, in 10 ms steps. We allowed 15 s between each pair of stimuli. Five responses were recorded at each inter-stimulus interval.

### Dentate spike analysis

Dentate spikes (DS) are intermittently occurring field events in the hilus caused by synchronous activation of the entorhinal input^6,7^ to DG. DS were identified in the hilus of DG as sharp, positive waves of LFP with prominence (distance between peak and closest preceding or following trough) above 2 S.D. of all detected positive peaks, and width between 2.5 and 12.5 ms measured at 50% of the peak’s prominence. Two types of DS – DS_L_ (type 1) and DS_M_ (type 2) were identified using a CSD fingerprinting method, where all DS that exhibited a symmetric pair of current sinks in the distal molecular layers of DG were identified as DS_L_, and DS that exhibited a symmetric pair of current sinks in the medial/proximal molecular layers of DG were identified as DS_M_.

### DREADD inactivation of the medial perforant path to verify electrode placement

We used a transgenic mouse (Ent-tTA; Mutant Mouse Resource & Research Centers, Stock: 031779-MU; RRID: MMRRC_031779-MU) bred to a hM4Di-tetO mouse (Jackson Laboratory). The resulting mouse can be used to suppress neurons in the superficial layers of medial entorhinal cortex (MEC) that give rise to the MPP ^8,9^. This transgenic mouse expressed the inhibitory DREADD receptor HM4Di in the MEC layers 2 and 3. CNO injection activated the hM4Di receptors that hyperpolarize the principal cells in superficial layers of MEC and in so doing, inactivated the MPP pathway. Prior to CNO administration, a baseline recording was performed and clear layer specific evoked responses in DG were elicited with test stimuli in the range from 0 to 250 μA. One hour after systemic i.p. injection of CNO, the evoked responses were recorded again. CNO produced a dose-dependent weakening of both the evoked dendritic responses and the evoked somatic responses, confirming targeting of the MPP-DG connection. The CNO effect was reversible which was confirmed by repeating the test stimuli 24 h after CNO administration, at which time the evoked responses were observed to return to the baseline levels, before CNO.

### In vivo acute surgery and recording

Subjects were 19 adult male and female Gad2Cre-ChR2eYFP mice weighing 25-30 g and aged 2-6 months. Gad2Cre-ChR2eYFP mice were bred by crossing a Gad2-IRES-Cre knock-in mouse (Jax stock number 028867 http://www.informatics.jax.org/accession/MGI:4418713 *Gad2^tm2(cre)Zjh^*) with a mouse that expresses a floxed channelrhodopsin-2/EYFP fusion protein (Jax stock number 012569 B6;129S-*Gt(ROSA)26Sor^tm32(CAG-COP4*H134R/EYFP)Hze^*/J) so it will express the fusion protein in Gad2+ cells (inhibitory neurons) expressing Cre recombinase. Mice were anesthetized with 1.20 g/kg urethane (i.p.) in three injections spaced 15 min apart, and placed in a stereotaxic frame. The mice were surgically implanted with recording and stimulating electrodes. The recording and stimulation system was identical to what was used for the freely-behaving mouse recordings, but there was an addition made to also enable optogenetic stimulation. For optogenetic stimulation, a 473 nm laser (OEM laser systems Midvale, UT) was attached to the implanted optic fibers. Peak laser power in the brain was 10 mW. After checking optic stimulation responses by changing laser power from 0 to 10mW in 1mW steps, the lowest light intensity to induce 90-100% of the maximum responses was used for optic stimulation.

### Experimental design for assessing CCT-induced changes in MPP-DG circuit function

Seven mice received CCT according to the identical initial training protocol that was used for behavioral assessment of learning to learn (Fig. 1). A 30-minute pretraining session was followed by 3 daily 30-min training trials with one 30-min retention session after 1 week. Four mice in the PL group received the same training protocol but with shallow water on the arena. Six mice in the SE control group received the same exposure to the environment but without shock. Mice in the home cage group (n = 5) received the same transfer to the behavior testing room but only stayed in their home cage during another mouse’s training session. Two hours after the retention test, the mice were anesthetized with 20% urethane as described above. Recording sessions consisted of 3 experiments. 1) Input-output (I-O) curve with electric MPP stimulation, 2) I-O curve with light stimulation in DG and electric stimulation in MPP, 3) a paired pulse inhibition experiment with MPP electrical stimulation. First, evoked responses in dorsal hippocampus to the MPP stimulation were measured by 100 ms pulse stimulation in the range 0-300 μA in 50 μA steps. Each stimulus was repeated 4 times. Next, a 5 ms light pulse in the DG was given 0 ms or 5 ms before the MPP stimulation. The evoked potential was measured as described before. Finally, paired pulse inhibition was measured. A stimulation intensity that induces 70% of the maximum evoked response was chosen for the PPI experiments. Two-MPP pulses were given in succession at varying intervals.

### Ex vivo acute hippocampus slice evoked potential recording

The subjects were 22 adult GadCre-ChR2-eYFP animals and 7 adult C57/BL6J mice, all mice were 25-30 g and aged 3-6 months. The behavioral training protocol was identical to the awake behaving and *in vivo* anesthetized experiments. One week after the last training day, retention of the conditioned avoidance was tested and the mice were returned to their home-cage for 1 hour. The mice were then transferred into an anesthesia induction chamber and kept there to habituate to the chamber for 10 min. The mice were then deeply anesthetized with isoflurane (5% in 100% oxygen) for 3 min, and immediately euthanized by rapid decapitation. Transverse slices (400 μm) from the right dorsal hippocampus were obtained for *ex vivo* electrophysiology experiments. Brain slices were cut with a vibratome (VT1000, Leica, Germany) in ice cold artificial cerebrospinal fluid (ACSF). The ACSF was comprised of the following solutions in mM concentrations: 119 NaCl, 4.0, KCl, 1.5 MgSO_4_, 2.5 CaCl_2_, 26.2 NaHCO_3_, 1 NaH_2_PO_4_ and 11 Glucose, saturated with 95% O2, 5% CO2) and then warmed in oxygenated ACSF to 35°C for 1 hour. The slices were allowed to equilibrate for at least 60 min in oxygenated ACSF at room temperature. For experiments, the slices were immersed in a recording chamber (Scientifica Inc., UK) with oxygenated ACSF at 35–36°C. Field excitatory postsynaptic potentials (fEPSP) from the termination zones of the MPP in the supraDG and infraDG were recorded in response to stimulation of MPP fibers with bipolar electrodes (FHC & Co, ME, USA). The stimulus pulse duration was 100 μs with 30 s between stimuli. The recording electrodes were borosilicate glass pipettes taper-pulled to an impedance of 5-7 MOhm (Sutter Inst. Co., Novato, CA, USA) and filled with the ACSF solution. To examine the persistent functional changes associated with CCT, we characterized synaptic transmission by the I-O curve and paired-pulse inhibition (PPI) protocols. Synaptic transmission measured as the stimulus-response relationship between the input stimulation voltage and slope of the fEPSP response were generated at increasing voltage stimulation. The fEPSPs at the supra- and infrapyramidal layers of DG were recorded separately. A 200-μm diameter optic fiber (Thorlabs Inc. NJ, USA) was connected to an optic cable to deliver the light stimulation to the hippocampus slices. The optic fiber was positioned as close to the recording electrode as practical, such that the distance between and recording electrode and optic fiber was less than 200 μm. The same 473 nm laser that was used for the anesthetized *in vivo* recordings, was used and 1 ms of 5 mW light was used for stimulation. Electrophysiological data from hippocampal slices were acquired with PClamp 11 software (Molecular Devices, Sunnyvale, CA, USA) and data analysis was performed with Clampfit (Molecular Devices, Sunnyvale, CA, USA).

### eYFP-labeling of GABAergic inhibitory neurons

Ten GadCre-eYFP mice were used. Ninety minutes after the last behavioral training session, the mice were anesthetized with a sodium pentobarbital and sodium phenytoin solution (EUTHASOL, Virbac AH, Inc. 20mg/ml) and transcardially perfused with 50 ml saline, followed by 50 ml ice-cold 4% paraformaldehyde (PFA) in saline. The brains were removed and post-fixed overnight in 4% PFA at 4°C. The brains were then cryoprotected in 30% sucrose for two days at 4°C and subsequently frozen in optimum cutting temperature (OCT) compound (Tissue Tek, Torrance, CA) until cryostat sectioning. Serial sections (40 μm) were cut through the entire dorsal hippocampus on a cryostat and stored in phosphate-buffered saline (PBS) with 0.02% NaN_3_, washed four times for 15 min in PBS and 5 min in PB, then mounted onto gelatin coated glass slides and cover slipped with a mountant (ProLong™ Glass Antifade Mountant, Thermofisher) for microscopic analysis. Two 20X images per side of dentate gyrus in the dorsal hippocampus and one 10X image focusing on the dorsal hippocampus for each animal were captured by a Leica TCS SP5 confocal microscope (Leica, Wetzlar, Germany). Quantification was performed using ImageJ software (US National Institutes of Health) with the operator blinded to the experimental conditions.

### Statistical Comparisons

All data were plotted in the main text figures, when appropriate using Tukey whisker plots, which marks outliers (“x”) ^10^. Outliers were determined by the ROUT method ^11^ with the maximum false discovery rate Q = 1 %. The only data that were excluded came from a single PL animal (beh14) because of a technical error on the first subsequent training session (Ts1).” Within subject comparisons were performed, when possible, using paired statistics such as Student’s paired t test. Comparisons amongst multiple factors or amongst more than two groups were performed using ANOVA, with repeated measures (RM) when the effect of a time series or sequence of measurement factors, such as a learning curve or set of stimulus intensities was being evaluated. The Greenhouse-Geisser correction was used when the data being compared by RM ANOVA violated the sphericity assumption as assessed by Mauchly’s Test of Sphericity. Tukey’s Honest significant difference (HSD) tests were used to evaluate all post-hoc pairwise comparisons when appropriate, and Sidak’s multiple comparisons were used when only a subset of pairwise comparisons were of interest. The test statistics and degrees of freedom are presented along with the p values and effect sizes for all comparisons. A value of p < 0.05 was considered significant. Statistical evaluations were performed and figures created using JMP Pro 14.1.0 (SAS, Cary, NC) and Prism 8 (GraphPad, San Diego, CA).

## REFERENCES

1 Thomas, C. & Baker, C. I. Teaching an adult brain new tricks: a critical review of evidence for training-dependent structural plasticity in humans. Neuroimage 73, 225–236, doi:10.1016/j.neuroimage.2012.03.069 (2013).

2 Thomas, C. & Baker, C. I. On evidence, biases and confounding factors: Response to commentaries. Neuroimage 73, 265–267, doi:10.1016/j.neuroimage.2012.11.009 (2013).

3 Zatorre, R. J., Fields, R. D. & Johansen-Berg, H. Plasticity in gray and white: neuroimaging changes in brain structure during learning. Nat Neurosci 15, 528–536, doi:10.1038/nn.3045 (2012).

4 Alonso, A., van der Meij, J., Tse, D. & Genzel, L. Naïve to expert: Considering the role of previous knowledge in memory. Brain and Neuroscience Advances 4, 2398212820948686, doi:10.1177/2398212820948686 (2020).

5 Bavelier, D., Green, C. S., Pouget, A. & Schrater, P. Brain plasticity through the life span: learning to learn and action video games. Annu Rev Neurosci 35, 391–416, doi:10.1146/annurev-neuro-060909-152832 (2012).

6 Bartlett, F. C. Remembering: A Study in Experimental and Social Psychology. (Cambridge University Press, 1932).

7 Harlow, H. F. The formation of learning sets. Psychol Rev 56, 51–65 (1949).

8 Yeager, D. S. et al. A national experiment reveals where a growth mindset improves achievement. Nature, doi:10.1038/s41586-019-1466-y (2019).

9 Brewin, C. R. Theoretical foundations of cognitive-behavior therapy for anxiety and depression. Annu Rev Psychol 47, 33–57, doi:10.1146/annurev.psych.47.1.33 (1996).

10 Anguera, J. A. et al. Video game training enhances cognitive control in older adults. Nature 501, 97–101, doi:nature12486 [pii] 10.1038/nature12486 (2013).

11 Subramaniam, K. et al. Computerized cognitive training restores neural activity within the reality monitoring network in schizophrenia. Neuron 73, 842–853, doi:S0896-6273(12)00049-9 [pii] 10.1016/j.neuron.2011.12.024 (2012).

12 Froemke, R. C. et al. Long-term modification of cortical synapses improves sensory perception. Nat Neurosci 16, 79–88, doi:10.1038/nn.3274 (2013).

13 de Villers-Sidani, E. et al. Recovery of functional and structural age-related changes in the rat primary auditory cortex with operant training. Proc Natl Acad Sci U S A 107, 13900–13905, doi:10.1073/pnas.1007885107 (2010).

14 Pavlowsky, A., Wallace, E., Fenton, A. A. & Alarcon, J. M. Persistent modifications of hippocampal synaptic function during remote spatial memory. Neurobiol Learn Mem 138, 182–197, doi:10.1016/j.nlm.2016.08.015 (2017).

15 Cimadevilla, J. M., Wesierska, M., Fenton, A. A. & Bures, J. Inactivating one hippocampus impairs avoidance of a stable room-defined place during dissociation of arena cues from room cues by rotation of the arena. Proc Natl Acad Sci U S A 98, 3531–3536, doi:10.1073/pnas.05162839898/6/3531 [pii] (2001).

16 Cimadevilla, J. M., Fenton, A. A. & Bures, J. Functional inactivation of dorsal hippocampus impairs active place avoidance in rats. Neurosci Lett 285, 53–56, doi:S0304-3940(00)01019-3 [pii] (2000).

17 Hsieh, C. et al. Persistent increases of PKMzeta in memory-activated neurons trace LTP maintenance during spatial long-term memory storage. Eur J Neurosci, doi:10.1111/ejn.15137 (2021).

18 Hsieh, C. et al. Persistent increased PKMzeta in long-term and remote spatial memory. Neurobiol Learn Mem 138, 135–144, doi:10.1016/j.nlm.2016.07.008 (2017).

19 Tsokas, P. et al. Compensation for PKMzeta in long-term potentiation and spatial long-term memory in mutant mice. Elife 5, e14846, doi:10.7554/eLife.14846 (2016).

20 Pastalkova, E. et al. Storage of spatial information by the maintenance mechanism of LTP. Science 313, 1141–1144, doi:10.1126/science.1128657 (2006).

21 van Dijk, M. T. & Fenton, A. A. On How the Dentate Gyrus Contributes to Memory Discrimination. Neuron 98, 832–845, doi:10.1016/j.neuron.2018.04.018 (2018).

22 Talbot, Z. N. et al. Normal CA1 Place Fields but Discoordinated Network Discharge in a Fmr1-Null Mouse Model of Fragile X Syndrome. Neuron 97, 684–697, doi:10.1016/j.neuron.2017.12.043 (2018).

23 Wesierska, M., Dockery, C. & Fenton, A. A. Beyond memory, navigation, and inhibition: behavioral evidence for hippocampus-dependent cognitive coordination in the rat. J Neurosci 25, 2413–2419, doi:25/9/2413 [pii] 10.1523/JNEUROSCI.3962-04.2005 (2005).

24 Dvorak, D., Radwan, B., Sparks, F. T., Talbot, Z. N. & Fenton, A. A. Control of recollection by slow gamma dominating mid-frequency gamma in hippocampus CA1. PLoS Biol 16, e2003354, doi:10.1371/journal.pbio.2003354 (2018).

25 Whishaw, I. Q. & Gorny, B. Path integration absent in scent-tracking fimbria-fornix rats: evidence for hippocampal involvement in “sense of direction” and “sense of distance” using self-movement cues. J Neurosci 19, 4662–4673 (1999).

26 Jezek, K. et al. Stress-Induced Out-of-Context Activation of Memory. PLoS Biol 8, e1000570, doi:10.1371/journal.pbio.1000570 (2010).

27 Garner, A. R. et al. Generation of a synthetic memory trace. Science 335, 1513–1516, doi:335/6075/1513 [pii] 10.1126/science.1214985 (2012).

28 Witter, M. P. The perforant path: projections from the entorhinal cortex to the dentate gyrus. Prog Brain Res 163, 43–61, doi:S0079-6123(07)63003-9 [pii] 10.1016/S0079-6123(07)63003-9 (2007).

29 O’Reilly, K. C., Perica, M. I. & Fenton, A. A. Synaptic plasticity/dysplasticity, process memory and item memory in rodent models of mental dysfunction. Schizophr Res, doi:10.1016/j.schres.2018.08.025 (2019).

30 Nitz, D. & McNaughton, B. Differential modulation of CA1 and dentate gyrus interneurons during exploration of novel environments. J Neurophysiol 91, 863–872, doi:10.1152/jn.00614.2003 (2004).

31 Madar, A. D., Ewell, L. A. & Jones, M. V. Temporal pattern separation in hippocampal neurons through multiplexed neural codes. PLOS Computational Biology 15, e1006932, doi:10.1371/journal.pcbi.1006932 (2019).

32 Ruediger, S. et al. Learning-related feedforward inhibitory connectivity growth required for memory precision. Nature 473, 514–518 (2011).

33 Scharfman, H. E., Sollas, A. L., Smith, K. L., Jackson, M. B. & Goodman, J. H. Structural and functional asymmetry in the normal and epileptic rat dentate gyrus. J Comp Neurol 454, 424–439, doi:10.1002/cne.10449 (2002).

34 Czeh, B., Abraham, H., Tahtakran, S., Houser, C. R. & Seress, L. Number and regional distribution of GAD65 mRNA-expressing interneurons in the rat hippocampal formation. Acta Biol Hung 64, 395–413, doi:10.1556/ABiol.64.2013.4.1 (2013).

35 Mercer, L. F., Jr., Remley, N. R. & Gilman, D. P. Effects of urethane on hippocampal unit activity in the rat. Brain Res Bull 3, 567–570, doi:10.1016/0361-9230(78)90089-8 (1978).

36 Shirasaka, Y. & Wasterlain, C. G. The effect of urethane anesthesia on evoked potentials in dentate gyrus. European Journal of Pharmacology 282, 11–17 (1995).

37 Olypher, A. V., Klement, D. & Fenton, A. A. Cognitive disorganization in hippocampus: a physiological model of the disorganization in psychosis. J Neurosci 26, 158–168, doi:26/1/158 [pii] 10.1523/JNEUROSCI.2064-05.2006 (2006).

38 Ferrante, M., Migliore, M. & Ascoli, G. A. Feed-forward inhibition as a buffer of the neuronal input-output relation. Proc Natl Acad Sci U S A 106, 18004–18009, doi:10.1073/pnas.0904784106 (2009).

39 Vasuta, C. et al. Metaplastic Regulation of CA1 Schaffer Collateral Pathway Plasticity by Hebbian MGluR1a-Mediated Plasticity at Excitatory Synapses onto SomatostatinExpressing Interneurons. eNeuro 2, doi:10.1523/ENEURO.0051-15.2015 (2015).

40 Donato, F., Rompani, S. B. & Caroni, P. Parvalbumin-expressing basket-cell network plasticity induced by experience regulates adult learning. Nature 504, 272–276, doi:10.1038/nature12866 (2013).

41 Letzkus, J. J., Wolff, S. B. & Luthi, A. Disinhibition, a Circuit Mechanism for Associative Learning and Memory. Neuron 88, 264–276, doi:10.1016/j.neuron.2015.09.024 (2015).

42 Ling, D. S. et al. Protein kinase Mzeta is necessary and sufficient for LTP maintenance. Nat Neurosci 5, 295–296, doi:10.1038/nn829 (2002).

43 Takeuchi, T., Duszkiewicz, A. J. & Morris, R. G. The synaptic plasticity and memory hypothesis: encoding, storage and persistence. Philos Trans R Soc Lond B Biol Sci 369, 20130288, doi:rstb.2013.0288 [pii] 10.1098/rstb.2013.0288 (2013).

44 Dvorak, D., Chung, A., Park, E. H. & Fenton, A. A. Dentate spikes and external control of hippocampal function. bioRxiv 2020.07.20.211615, doi:10.1101/2020.07.20.211615 (preprint).

45 Lee, J. W. & Jung, M. W. Separation or binding? Role of the dentate gyrus in hippocampal mnemonic processing. Neurosci Biobehav Rev 75, 183–194, doi:10.1016/j.neubiorev.2017.01.049 (2017).

46 Lee, J. W., Kim, W. R., Sun, W. & Jung, M. W. Role of dentate gyrus in aligning internal spatial map to external landmark. Learn Mem 16, 530–536, doi:10.1101/lm.1483709 (2009).

47 Luna, V. M. et al. Adult-born hippocampal neurons bidirectionally modulate entorhinal inputs into the dentate gyrus. Science 364, 578–583, doi:10.1126/science.aat8789 (2019).

48 Lee, H. et al. Early cognitive experience prevents adult deficits in a neurodevelopmental schizophrenia model. Neuron 75, 714–724, doi:S0896-6273(12)00578-8 [pii] 10.1016/j.neuron.2012.06.016 (2012).

## SUPPLEMENTARY REFERENCES

1 Franklin, K. B. J. & Paxinos, G. The mouse brain in stereotaxic coordinates. 3rd edn, (Academic Press, 2007).

2 Harrison, R. R. et al. A Low-Power Integrated Circuit for a Wireless 100-Electrode Neural Recording System. IEEE Journal of Solid-State Circuits 42, 123–133, doi:10.1109/JSSC.2006.886567 (2007).

3 Hrabetova, S. Extracellular diffusion is fast and isotropic in the stratum radiatum of hippocampal CA1 region in rat brain slices. Hippocampus 15, 441–450, doi:10.1002/hipo.20068 (2005).

4 Holsheimer, J. Electrical conductivity of the hippocampal CA1 layers and application to current-source-density analysis. Exp Brain Res 67, 402–410 (1987).

5 Pettersen, K. H., Devor, A., Ulbert, I., Dale, A. M. & Einevoll, G. T. Current-source density estimation based on inversion of electrostatic forward solution: effects of finite extent of neuronal activity and conductivity discontinuities. J Neurosci Methods 154, 116–133, doi:10.1016/j.jneumeth.2005.12.005 (2006).

6 Penttonen, M., Kamondi, A., Sik, A., Acsady, L. & Buzsaki, G. Feed-forward and feedback activation of the dentate gyrus in vivo during dentate spikes and sharp wave bursts. Hippocampus 7, 437–450, doi:10.1002/(SICI)1098-1063(1997)7:4<437::AID-HIPO9>3.0.CO;2-F (1997).

7 Bragin, A., Jando, G., Nadasdy, Z., van Landeghem, M. & Buzsaki, G. Dentate EEG spikes and associated interneuronal population bursts in the hippocampal hilar region of the rat. J Neurophysiol 73, 1691–1705 (1995).

8 Garner, A. R. et al. Generation of a synthetic memory trace. Science 335, 1513–1516, doi:335/6075/1513 [pii] 10.1126/science.1214985 (2012).

9 Kanter, B. R. et al. A Novel Mechanism for the Grid-to-Place Cell Transformation Revealed by Transgenic Depolarization of Medial Entorhinal Cortex Layer II. Neuron 93, 1480–1492 e1486, doi:10.1016/j.neuron.2017.03.001 (2017).

10 Krzywinski, M. & Altman, N. Visualizing samples with box plots. Nat Methods 11, 119–120, doi:10.1038/nmeth.2813 (2014).

11 Motulsky, H. J. & Brown, R. E. Detecting outliers when fitting data with nonlinear regression – a new method based on robust nonlinear regression and the false discovery rate. BMC Bioinformatics 7, 123, doi:10.1186/1471-2105-7-123 (2006).

